# The TRP-channel *painless* mediates substrate stiffness sensing in the legs during *Drosophila* oviposition

**DOI:** 10.1101/2025.08.18.670869

**Authors:** Vijayaditya Ray, Lasse B. Bräcker, Alexandros Kourtidis, Charlie Rosher, Gesa F. Dinges, Anna Pierzchlińska, Ansgar Büschges, Kai Feng, Kevin M. Cury, Nicolas Gompel

## Abstract

The distinct textural properties of fruits in varying stages of ripening present unique ecological opportunities for several species of fruit flies, resulting, over evolutionary times, in specialized egg-laying behaviors. In this study we identified a TrpA channel-dependent mechanosensory pathway in the legs, through the gene *painless,* that modulates the discernment of softer patches for oviposition in gravid *D. melanogaster* females. We report that the stiffness-sensing role of tarsi is mediated through external sensory organs housed, namely ventral mechanosensory bristles and subsets of campaniform sensilla present primarily at the joints between tarsomeres. Our findings provide new evidence that campaniform sensilla function as indirect stiffness sensors of oviposition substrates, owing to their placement at joints that experience maximal cuticular distortion. We show that Painless is expressed in mechanosensory neurons innervating peripheral organs where it likely participates in the transduction of stiffness-evoked stimuli. Furthermore, we observed that overexpression of *painless* in both campaniform sensilla and mechanosensory bristles partially rescues preference for the softer substrates in *painless* mutants, indicating that *painless* activity in these organs is necessary to mediate the preference. We propose that different interactions with a soft *vs.* a hard substrate (compression of the cuticle, distribution of contacts) results in differential mechanotransduction in *painless*-expressing neurons, determining oviposition preferences.

## Introduction

In oviparous animals that do not brood, such as fruit flies, choosing a suitable environment to deposit their eggs is critical to the survival of their progeny ^1^. It is well established that gravid female flies thoroughly evaluate the properties of the substrate onto which they will deposit their eggs, and that their choice is in line with the quality and benefits of resources for their progeny ^2^. This process of committing to a particular site relies on the reception and integration of various environmental sensory cues ^2–7^, as well as on the mating status of the female ^8^. A female fruit fly first employs long-range sensory modalities, vision and olfaction, to identify and navigate to a potential egg-laying site, before assessing its quality at short-range by direct contact, through gustation and mechanosensation ^7^. The contributions of chemosensory modalities such as olfaction ^1,3,9^ and gustation ^10^ in influencing oviposition decisions have been extensively studied both behaviorally and mechanistically. By contrast, the neurogenetic architecture that mediates perception of mechanical properties, such as substrate stiffness during oviposition site selection is less well understood. Mechanosensation is a significant sensory modality employed by fruit flies to evaluate stiffness and detect variation in textural properties of substrates ^11,12^. In feeding behaviors, specific quotients of hardness are representative of the quality of the food consumed by the flies, thus heavily influencing feeding preferences ^13,14^. In oviposition, fruit flies breed upon a variety of host substrates and demonstrate marked species-specific preferences for substrate stiffness corresponding to their native hosts. For instance, *D. suzukii* females prefer to oviposit in ripe fruits and similarly stiff agarose substrates, whereas *D. melanogaster* females choose to deposit eggs in relatively soft, decaying fruits, and similarly choose soft agarose substrates ^11^. Our aim with the present work was to understand the mechanisms underlying the assessment of a substrate’s physical properties in the context of egg laying. We have analyzed the mechanosensory system used by gravid *D. melanogaster* females during the selection of an egg-laying surface. We have identified and characterized stiffness sensing across multiple scales, from body appendages to sensory organs, and down to the level of neurons and genes. Our results show that females flies rely on information derived from campaniform sensilla and mechanosensory bristles of their tarsi (the distal-most segments of each leg) to choose an egg-deposition site and use the mechanoreceptor *painless* in this process.

## Results

### Tarsi are required to sense stiffness of oviposition substrates

We have previously shown that in a two-choice oviposition assay, gravid *D. melanogaster* females consistently show pronounced oviposition preferences for the softer option (0.25% agarose vs. x >0.25% agarose) ^11^. We therefore first investigated which appendages are involved in this choice. We targeted body parts engaged in direct contact with a potential egg-laying substrate prior to oviposition ^15,16^. In an initial assay, we ablated either the distal mouthparts or the tarsi from mated females and assessed their ability to choose between substrates of different stiffness. Since the ovipositor is engaged prior to and throughout oviposition ^16^, removing this organ was not an option. In a two-choice group assay, offering soft and hard substrate options (0.25% agarose and 0.75% agarose, corresponding to the stiffness of rotten and ripe fruits respectively ^11^; Fig. 1A), groups of 40 experimental flies with ablated appendages or control flies with intact appendages were introduced into the chamber and allowed to lay eggs over a period of 16 hours. The preference for depositing eggs on the softer substrate was then calculated. We observed that proboscis ablation did not cause any impairment in oviposition preference for the softer substrate compared to control flies with an intact proboscis (Fig. 1B, Fig. S1A). By contrast, tarsal ablation substantially reduced the preference for the softer substrate (Fig. 1C, Fig. S1B). Of note, flies with ablated appendages laid much fewer eggs than control flies, but enough to test their ability to compare substrates (Fig. S1A-B). These results suggest that tarsi are required to gauge the textural properties of potential egg-laying substrates.

**Figure 1.**
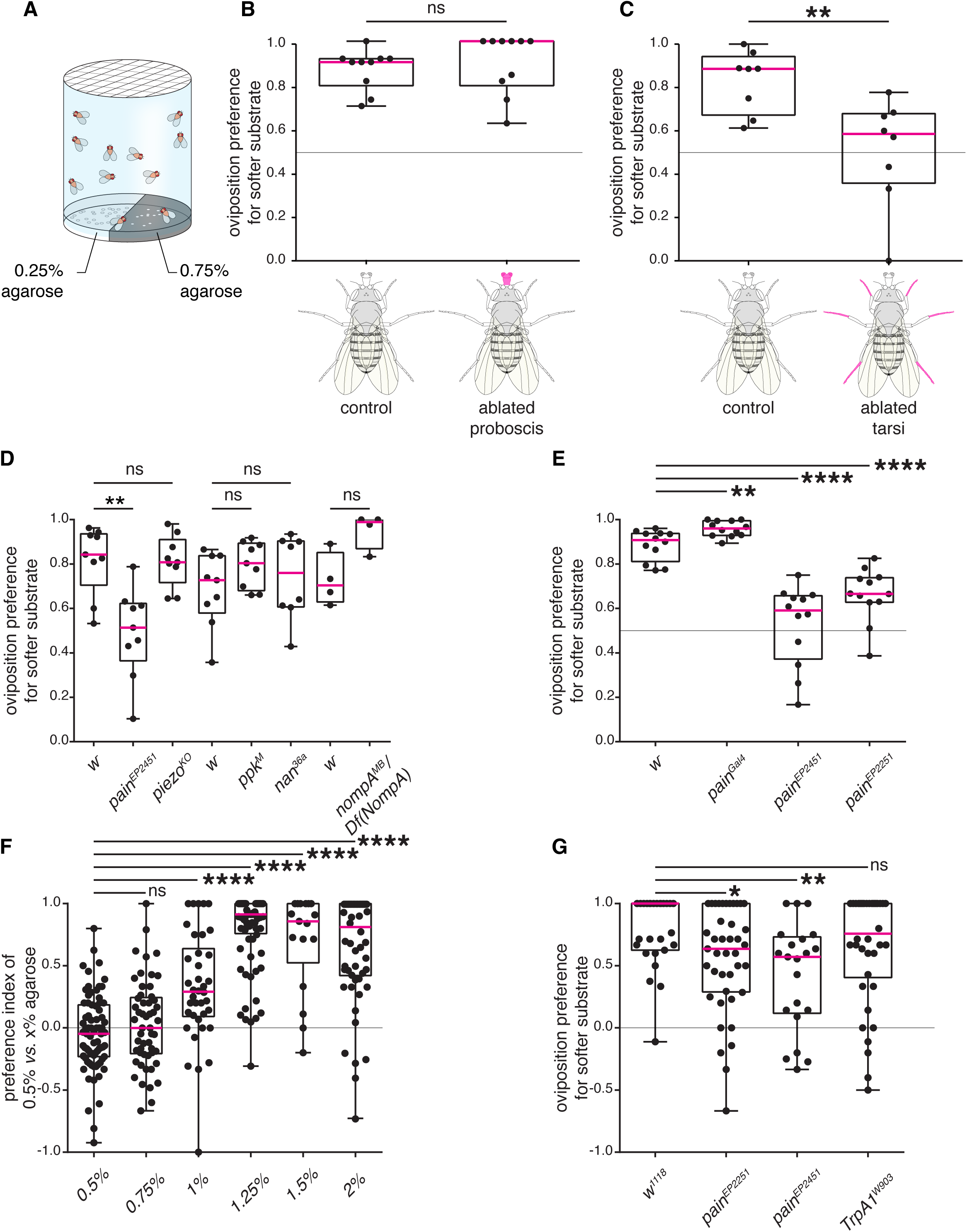
Female *Drosophila* use their tarsi and the gene *painless* to evaluate substrate stiffness during oviposition. (A) Schematic of the two-choice oviposition assay for substrate stiffness. Groups of ∼100-150 flies of mixed sexes can choose between 0.25% and 0.75% agarose supplemented with 0.5 M glucose, to lay their eggs. From the proportion of eggs laid on each side of the plate after 16 hours, we derived a preference index (PI) for the softer side ((eggs on 0.25% – eggs on 0.75%)/total number of eggs). (B-C) Ablation experiments, in which the proboscis (B, n=12) or all tarsi (C, n=8)) were clipped from females and their substrate preference were compared to those of control females with intact appendages. (D-E) Evaluating substrate stiffness preferences of mutants for known mechanoreceptors (D, n=9, 9, 9, 9, 9, 8, 4, and 4 respectively) pointed to a potential involvement of the gene *painless*, for which we re-tested additional *pain* alleles in group assays (E, n=12, 12, 12, and 12, respectively). (F) Testing individual wild-type females in a two-choice single fly assay with increasing difference in agarose concentration between the soft and the hard substrates (agarose concentration increasing from 0.5%-2%; n=75, 53, 38, 76, 17, and 54, respectively). (G) Testing mutants of TrpA channels for their oviposition preferences for the softer side (0.5%) in a two-choice single fly assay (n=27, 41, 21, and 38). In (B-E), group two-choice stiffness assays (0.25% vs 0.75% agarose) were used and each dot on the graphs represents the preference index of a single replicate involving a batch of females. In these assays, the agarose substrates were supplemented with 0.5 M. In (F-G), single-fly two-choice stiffness assays were used in which females can individually choose between 0.5% and 1.25% agarose, supplemented with only 3% acetic acid (F) or 0.5 M glucose and 3% acetic acid (G). Each dot in figures (F-G) represents a separate mated female of a particular genotype. In all figures, the median of computed PIs are depicted in magenta within the box and the whiskers extend to the true minimum and maximum values for each genotype. p values for statistical significance were calculated using a non-parametric Mann Whitney t-test. ns, non-significant, p > 0.05; *, p < 0.05; **, p < 0.01; ***, p < 0.005; ****, p < 0.0001. See Fig. S1 for total egg counts.

### The TrpA channel *painless* functions as a mechanoreceptor that senses stiffness of oviposition substrates

The tarsi harbor a variety of mechanosensory organs ^17,18^ and express a host of proteins involved in mechanotransduction ^12^. We first asked which mechanoreceptor could potentially sense the stiffness of a potential oviposition surface by screening available mutants for several genes with reported functions in mechanosensory signal reception or transduction pathway ^12,19–22^. Specifically, we tested whether females mutant for *nan*, *NompA*, *painless*, *piezo* and *ppk* displayed altered oviposition preferences for softer surfaces compared to controls. Using a two-choice group assay (0.25% *vs.* 0.75% agarose), we observed that mutants for the genes *NompA, piezo, ppk,* and *nan* showed similar oviposition preferences as control flies (Fig. 1D, Fig. S1C). However, flies homozygous for different alleles of the gene *painless* (*pain^EP2251^* and *pain^EP2451^*) ^23,24^, showed a prominent and statistically significant reduction in their preference to deposit eggs on the softer substrate (Fig. 1D-1E, Fig. S1C-1D). We also tested the *pain-Gal4* allele that contains a Gal4 insertion in the first exon of the *painless* locus and has been previously reported to show intermediate isothiocyanate avoidance defects ^25^. Surprisingly, *pain-Gal4* flies showed mild but significantly higher oviposition preferences for the softer side.

To further confirm the phenotype associated with *pain* with more alleles, we developed a high-throughput assay to determine the preference of individual gravid females. In a custom-made behavioral apparatus, which simultaneously tests 56 flies (see Methods), each fly was exposed to two adjacent substrate patches within a small cubical chamber. To examine baseline stiffness preferences of wild-type flies in this assay, we first tested a range of choices, maintaining 0.5% agarose concentration constant as the softer side, and incrementally increasing the harder side by 0.25% points. We observed that the oviposition preference for the softer side increased until the harder side reached 1.25% agarose and then plateaued for concentrations above 1.25% (Fig. 1F, Fig. S1E). We therefore selected 0.5% *vs.* 1.25% agarose for our subsequent single-fly two-choice stiffness assays.

We confirmed that in this single-fly behavioral set-up, homozygous *pain* mutants also exhibit a deficit in oviposition preference for the softer side, compared to control flies (Fig. 1G, Fig. S1F). To further test the specificity of *pain* in this context, we tested a null mutant allele of another *TRPA* family member, *TrpA1* ^26^, whose function was shown to overlap with that of *pain* in response to horseradish/wasabi derived allyl isothiocyanate ^27^. Mutant *TrpA1^W903^*females showed preference for the softer substrate, like control flies (Fig. 1G, Fig. S1F). Altogether, the results strongly posit *pain* as a specific mechanoreceptor necessary for sensing stiffness of egg-laying substrates.

### *painless*-expressing neurons of the leg mediate the choice of a softer egg-laying substrate

The comparable impairment to select softer substrates upon tarsal ablation and *pain* mutation (Figs. 1C, 1E, 1G) suggested to us that *pain* may be functional in the tarsi. We hypothesized that *pain*-expressing neurons innervating peripheral sensory organs of the legs may be required to select softer oviposition substrates. To test our hypothesis, we conditionally blocked neurotransmission in all *pain*-expressing neurons by expressing a thermosensitive allele of the exocytose gene *shibire* (*UAS-shi^ts^*) ^28^, under the control of *pain-Gal4,* which has been reported to label *pain*-positive sensory neurons in several parts of the body including the tips of the proboscis, inner pharynx, upper margins of the wings and the legs ^25^. Experimental flies in which synaptic transmission was blocked (at 31°C) in all *pain*-expressing neurons (*pain-Gal4>UAS-shi ^ts^*) showed compromised ability in choosing the softer substrate as compared to control flies in group assays (Fig. 2A, Fig. S2A). We concluded that *pain*-positive neurons are necessary in the choice of softer substrates. Since *pain-Gal4* expression is not limited to the legs, we could not exclude that the effect we observed resulted from the inactivation of *pain*-positive neurons elsewhere than in the legs. We therefore resorted to an intersectional genetic approach to selectively silence *pain*-expressing neurons innervating the legs by the targeted expression of the inward-rectifying potassium (K⁺) channel Kir2.1, which blocks neurotransmission by hyperpolarizing the neuron. We combined a *dac[H1]*-flippase line, driving flippase expression in the legs ^29^, to a conditional *UAS-FRT-stop-FRT Kir2.1* effector ^30^, containing a translational stop removable upon the action of the flippase, crossed with the *pain-Gal4* line. To verify the efficacy of the intersectional strategy, we first imaged the brain and the ventral nerve cord of the experimental flies to visualize the expression of the *UAS>myrtdTomato-SV40>eGFPKir2.1* cassette (Fig. S4A-A’). In this construct, tdTomato is expressed in *pain-Gal4*-positive cells in the absence of *dac-flp*, while Kir2.1::eGFP expression is restricted to cells where both *dac-flp* and *pain-Gal4* are active due to excision of the STOP cassette flanked by FRT sites. We observed Kir2.1::eGFP expression in many but not all leg-neuronal projections of the thoracic neuromeres (Fig. S4A’). The brains of these animals showed extremely sparse GFP expression in neuronal cell bodies (Fig. S4A) that was variable across samples (n=2). These flies with silenced *pain*-positive neurons in the legs (*pain-Gal4/+; dac[H1] Flp, UAS>stop>Kir2.1/+*) showed a pronounced reduction in their ability to select the softer surfaces compared to control flies when tested in the two-choice single fly oviposition assay (0.5% vs. 1.25% agarose as soft and hard substrates, respectively; Fig. 2B, Fig. S2B). These results are similar to the phenotype observed upon complete *pain-Gal4* silencing, thus reinforcing our hypothesis that *pain* is functioning in the legs to mediate the choice of an oviposition substrate.

**Figure 2.**
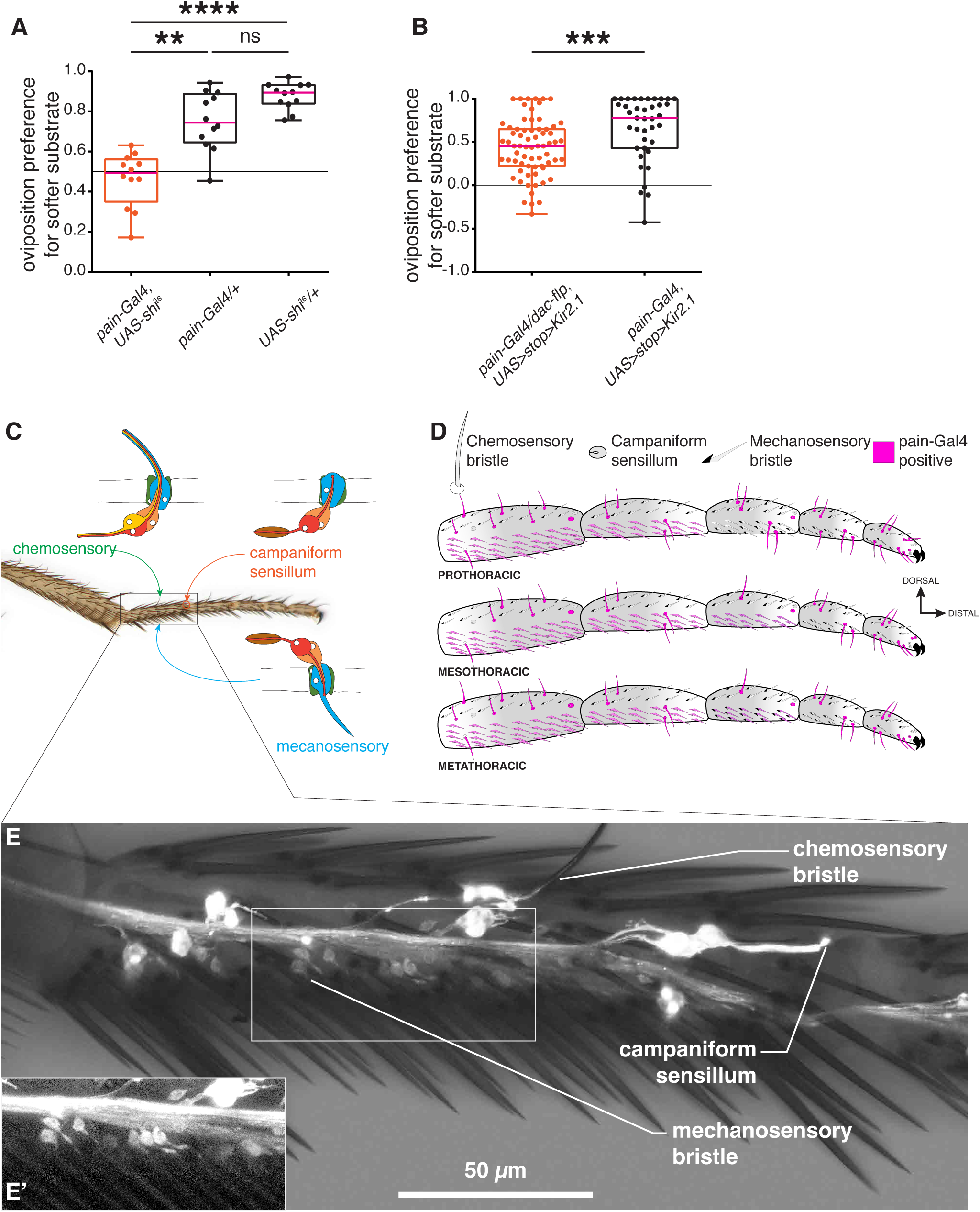
*pain* is expressed in neurons innervating chemosensory bristles, campaniform sensilla and mechanosensory bristles in the tarsi. (A-B) Global (A) or leg-restricted (B) blocking of neurotransmission in *pain*-positive neurons alters female oviposition preference on softer substrates.(A) a two-choice group stiffness assay (0.25% vs. 0.75% agarose) of *pain-Gal4, UAS-shi^ts^* females tested at 31°C (restrictive temperature, silenced, in red). Control animals (in black) only carry one allele of either *UAS-shi^ts^* or *pain-Gal4* (n=12,12, and 12, respectively). (B) Females with conditional inactivation of *pain-Gal4* neurons in the legs, (*pain-Gal4*, *dac-flippase*, *UAS>stop>Kir2.1*) tested in single-fly two-choice stiffness assay (0.5% vs. 1.25% agarose). Control animals lack *dac-flippase* (n=68 and 39, respectively). p values were calculated using Kruskal–Wallis test followed by a Dunn’s multiple comparisons test in (A) and a non-parametric Mann Whitney t-test in (B). ns, non-significant, p > 0.05; *, p < 0.05; **, p < 0.01; ***, p < 0.005; ****, p < 0.0001. See Fig. S2 for total egg counts. In this and all subsequent figures, experimental data points are in red, controls are in black. (C) Sensory organs of the tarsus (D) Schematic representation of *pain* expression in the tarsi. Sensory organs innervated by *pain*-positive neurons are in magenta. (E) *pain* expression in tarsomere 1 neurons of a prothoracic leg (composite image: brightfield and GFP): 3–4 neurons per chemosensory bristles, a campaniform sensillum neuron (1/2 of Ta1G ^31^) at the joint between tarsomeres Ta1 and Ta2, and single neurons for each ventral mechanosensory bristles (inset (E’)).

### *painless*-expressing neurons innervate campaniform sensilla, chemosensory bristles and mechanosensory bristles in the tarsi

Tarsi contain three types of external sense organs: campaniform sensilla ^31^, chemosensory bristles, and mechanosensory bristles ^17,18,32^. Previously, *pain* expression was reported in tarsus gustatory neurons ^25^, with no further details on their number and distribution. We examined *pain* expression in the tarsi to determine the identity and distribution of sense organs innervated by *pain*-expressing neurons. We imaged the dissected tarsi of *pain-Gal4, UAS-GFP* flies and observed that *pain-*expressing sensory neurons innervated all three types of external sense organs (Fig. 2C-E, Table S1). Campaniform sensilla (CS) in the tarsi are organized either as singlets in tarsomere 1(Ta1S) or in clusters of 2-4 sensilla in tarsomeres 1, 3 and 5 (respectively named Ta1G, Ta3G and Ta5G ^31^). *pain*-*Gal4* labelled neurons that innervate subsets of these CS, mostly positioned dorsally at the joints between tarsomeres or near the claws in the most distal tarsomere (Fig. 2D-E, Table S1). Furthermore, we observed *pain-Gal4>GFP* expression in neurons innervating chemosensory bristles (Fig. 2D-E, Table S1). These neurons, in bundles of 3-4 cells per organ, had their dendrites extending in the shaft of the bristle, a signature of chemosensory neurons ^33^. All chemosensory bristles in all legs showed positive GFP expression except for one in tarsomere 4 of the prothoracic legs. Chemosensory bristles are also innervated by a single mechanosensory neuron whose dendrite ends at the base of the shaft ^33^. We could not distinguish whether this neuron was also labelled. Finally, we observed GFP expression in neurons that innervate the ventrally positioned mechanosensory bristles of the tarsus (Fig. 2D-E’, Table S1). The expression was confined to all ventral mechanosensory organs of the proximal tarsomeres 1–2 in prothoracic legs, and 1–3 in meso- and metathoracic legs.

### Dorsal campaniform sensilla and ventral mechanosensory bristles of the legs gauge stiffness of egg-laying surfaces

Given the expression pattern of *pain* within the tarsi, we sought to uncover the distinct contribution of each type of external sense organs to stiffness preference. We leveraged previously identified Gal4 driver lines with specific expression in neurons innervating each organ type to manipulate their activity. *poxn-Gal4* expresses in chemosensory neurons innervating chemosensory bristles ^34^, *R38B08-Gal4* expresses in mechanosensory neurons innervating bristles ^35^. To target different and partially overlapping subsets of campaniform sensilla in the legs, we used the generation-1 driver line *R34E04-GAL4* (*CS1*) ^36^, as well as two split-GAL4 lines, *SS69574-Gal4* (*CS2*), and *SS69602-Gal4* (*CS3*) that we built (Fig. S3B-L). To disrupt neurotransmission in the sense organs labelled by these drivers, we either used *UAS-Kir2.1* or, in case of developmental lethality *(CS1* and *poxn-Gal4*), *UAS-shi^ts^*.

#### Silencing of chemosensory neurons

Expressing *UAS-shi^ts^* with *poxn-Gal4* at the restrictive temperature (31°C) resulted in significantly lower preferences for the softer patch when compared to the same genotype at the permissive temperature (21°C). However, we observed a similar effect for both controls (Gal4 alone and UAS alone; Fig. 3A, Fig. S5A). This result is inconclusive but suggests that the shift in preference is merely due to the change of temperature. *poxn-Gal4* is, however, not restricted to the legs ^34^, also driving expression in other appendages and the central nervous system. For a targeted neuronal inactivation of leg-specific neurons, we again utilized the intersectional strategy described above, combining *poxn-Gal4* with *dac-flp* to drive *UAS-FRT>stop>FRT Kir2.1*. We first validated it by imaging the brain and the ventral nerve cord to visualize the expression of the *UAS>myrtdTomato-SV40>eGFPKir2.1* cassette under the influence of *poxn-Gal4* driver. Flies in which Kir2.1::eGFP expression was restricted to the leg chemosensory neurons (*poxn-Gal4*, *dac-flp-UAS>stop>Kir2.1*) showed very sparse GFP expression in the brain (Fig. S4B). Likewise, in the ventral nerve cord, only a small subset of leg-specific neurons showed Kir2.1::eGFP expression in the thoracic neuromeres, while the rest expressed tdTomato (Fig. S4B’). Individual flies with silenced leg chemosensory neurons tested in a two-choice stiffness assay showed comparable oviposition preference to that of control flies (Fig. 3B, Fig. S5B). Because only a subset of the leg chemosensory neurons was silenced, however, as our confocal images showed, we cannot exclude a contribution of chemosensory neurons in stiffness sensing.

**Figure 3.**
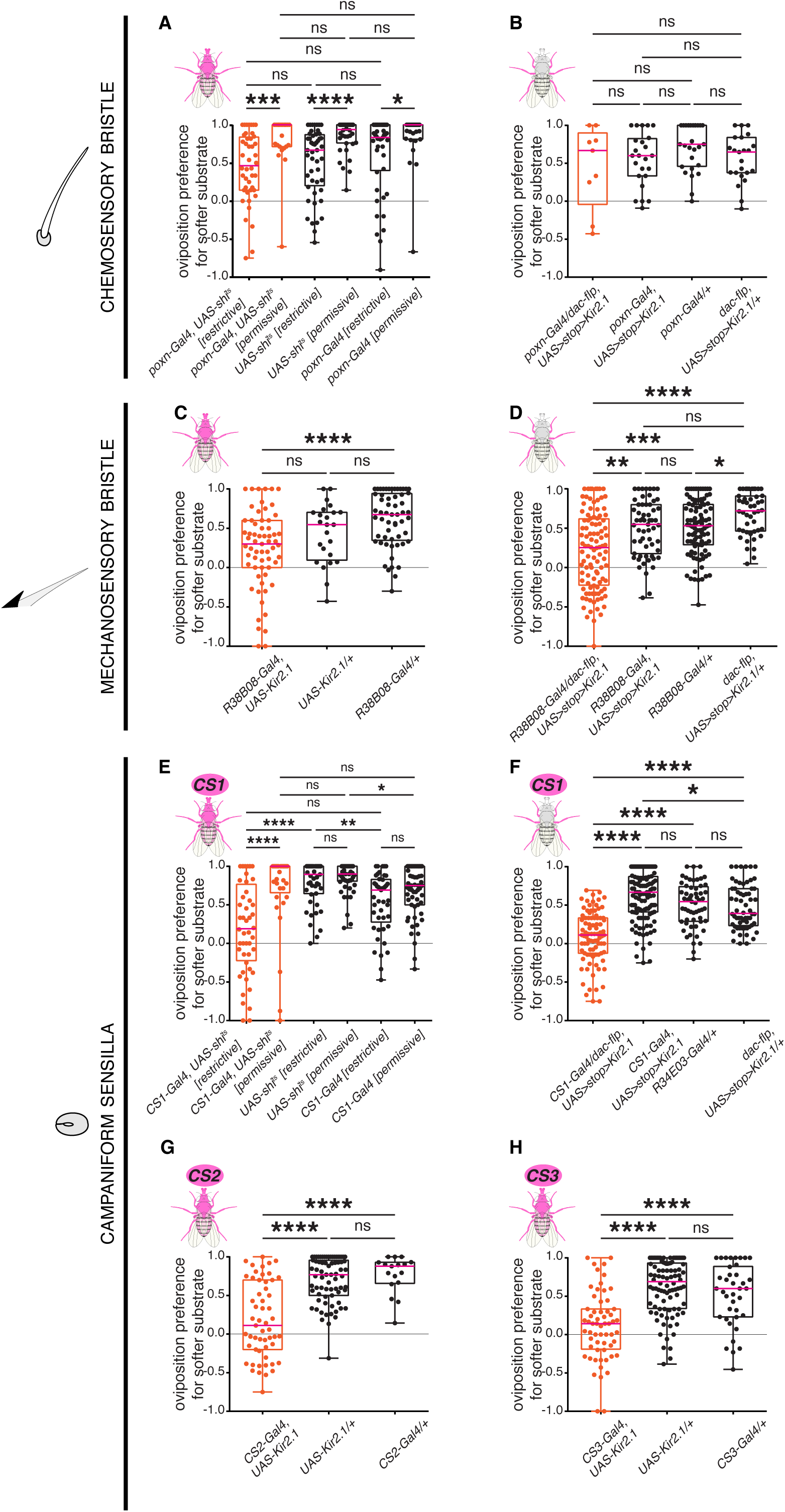
Leg campaniform sensilla and mechanosensory bristles mediate stiffness sensing of oviposition substrates. Single-fly two-choice stiffness assay (0.5% vs. 1.25% agarose) of females with global (A, C, E, G, H) or leg-restricted (B, D, F) blocking of neurotransmission in sensory neurons innervating chemosensory bristles (A-B), mechanosensory bristles (C-D) or campaniform sensilla (E-H). (A) *poxn-Gal4*, *UAS-shi^ts^* females at restrictive (silenced, 31°C) or permissive (non-silenced, 21°C) temperature, compared to *UAS*-alone or *Gal4*-alone controls. (B) conditional inactivation of *poxn-Gal4* neurons in the legs, (*poxn-Gal4*, *dac-flippase*, *UAS>stop>Kir2.1*), compared to controls lacking one of the genetic components (A: n=44, 23, 48, 38, 45, and 28, respectively; B: n=9, 24, 28, and 23, respectively). (C) *R38B08-Gal4*, *UAS-Kir2.1* females, compared to *UAS*-alone or *Gal4*-alone controls. (D) conditional inactivation of *R38B08-Gal4* neurons in the legs, (*R38B08-Gal4*, *dac-flippase*, *UAS>stop>Kir2.1*), compared to controls lacking one of the genetic components (C: n=63, 24, and 57, respectively; D: n=104, 59, 98, and 46, respectively). (E) *CS1-Gal4*, *UAS-shi^ts^*females at restrictive (silenced, 31°C) or permissive (non-silenced, 21°C) temperature, compared to *UAS*-alone or *Gal4*-alone controls. (F) conditional inactivation of *CS1-Gal4* neurons in the legs, (*CS1-Gal4*, *dac-flippase*, *UAS>stop>Kir2.1*), compared to controls lacking one of the genetic components (E: n=48, 33, 47, 52, 45, and 57, respectively; G: n=88, 121, 57, and 62, respectively). (G) *CS2-Gal4*, *UAS-Kir2.1* females, compared to *UAS*-alone or *Gal4*-alone controls. (H) *CS3-Gal4*, *UAS-Kir2.1* females, compared to *UAS*-alone or *Gal4*-alone controls (G: n=55, 71, and 17, respectively; H: n= 60, 92, and 38, respectively). p values for statistical significance were calculated using Kruskal–Wallis test followed by a Dunn’s multiple comparisons test. ns, non-significant, p > 0.05; *, p < 0.05; **, p < 0.01; ***, p < 0.005; ****, p < 0.0001. See Figs S3-S5 for expression patterns of driver lines, neurons labelled due to conditional inactivation and total egg counts.

#### Silencing of bristles mechanosensory neurons

Expressing *UAS-Kir2.1* using the complete expression profile of *R38B08-Gal4* resulted in a reduction in oviposition preferences for the softer substrate, which was only significant when compared to the Gal4 control (Fig. 3C, Fig. S5C). However, a trend of reduction of softer substrate preference was seen in *R38B08-Gal4*-silenced flies in comparison to both the controls. Since *R38B08-Gal4* is also not specific to the legs (FlyLight Project at Janelia Research Campus, HHMI ^36^), we used the dac-flippase mediated intersectional approach described above. Flies in which Kir2.1::eGFP expression was restricted to leg neurons (*R38B08-Gal4, dac-flp-UAS>stop>Kir2.1*) showed no eGFP expression in the brain (Fig. S4C) and labelled a subset of leg neuronal efferents in the ventral nerve cord (Fig. S4C’). In a single-fly assay, blocking neural transmission in these leg neurons significantly reduced egg-laying preferences for the softer side when compared to control flies (Fig. 3D, Fig. S5D), indicating a clear involvement of these neurons in the choice.

#### Silencing of campaniform sensilla mechanosensory neurons

To target campaniform sensilla, we first employed the broad driver *CS1-Gal4* that labels mechanosensory neurons innervating numerous CS in addition to some mechanosensory bristles in the legs. Driving *UAS-shi^ts^*with *CS1-Gal4* showed a significantly reduced preference in comparison to only the UAS control at the restrictive temperature (31°C) but a trend of reduction was observed with respect to both controls (Fig. 3E, Fig. S5E). At the permissive temperature (21°C), however, *CS1-Gal4*>*UAS-shi^ts^*behaves similarly to the controls (Fig. 3E, Fig. S5E). Since *CS1-Gal4* is not restricted to the legs, also showing expression in the central nervous system, we again used *dac-flippase* to limit Kir2.1-mediated silencing to leg-specific neurons. Leg-restricted *CS1-Gal4*-silenced flies showed no Kir2.1::eGFP expression in the brain (Fig. S4D) and showed positive expression in a subset of leg neuron projections in the ventral nerve cord (Fig. S4D’). When these *CS1-Gal4⋂dac*-silenced flies were tested in a single-fly two-choice assay, they showed significantly lower oviposition preferences for the softer substrate compared to the controls (Fig. 3F, Fig. S5F). These results indicate a role of a subset of mechanosensory neurons associated with both mechanosensory bristles and CS in the legs in preferences for the softer substrate. To isolate the specific contribution of campaniform sensilla, we generated two split-Gal4 drivers strictly restricted to these organs, CS2 and CS3, already referred to above. Both show peripheral projections but no cell bodies in the central nervous system (Figs. S4E-E’, S4F-F’). Driving *UAS-Kir2.1* with both *CS2-Gal4* and *CS3-Gal4* (*CS-Gal4*, *UAS-Kir2.1*) resulted in significantly reduced oviposition preference for the softer side compared to control flies (*CS-Gal4* alone and *UAS-Kir2.1* alone; Figs. 3G-H, S5G-H). To rule out the possibility that the preference phenotype results from locomotor deficits compromising female choices upon CS or mechanosensory bristle neuron inactivation, we tested the different Gal4 lines in an oviposition assay focusing on substrate chemical composition rather than stiffness. Here, females were offered a substrate with 1% agarose, with one side only supplemented with 200 mM glucose. We hypothesized that inactivating mechanosensory neurons of the legs should not affect female chemosensory perception, and therefore, any shift in egg-laying preference compared to genetic controls would be attributed to locomotor defects. We observed that blocking neurotransmission with *UAS-Kir2.1* in all neurons labelled by *CS2-Gal4* and *CS3-Gal4* did not significantly affect oviposition preference for sugar-containing substrates in comparison to the Gal4 control. The UAS control surprisingly showed significantly higher preferences for the sugar-containing substrate compared to both the experimental genotype and the Gal4 control (Fig. S6A-B, Fig. S7A-B). Expressing Kir2.1 in all bristle mechanosensory neurons labelled by *R38B08-Gal4* and leg-specific mechanosensory neurons labelled by *CS1-Gal4* resulted in comparable oviposition preference for sugar-containing substrates as both the genetic controls (Fig. S6C-D, Fig. S7C-D). These results clearly indicate that flies with silenced mechanosensory neurons are not compromised in their ability to express oviposition stiffness preferences due to locomotor defects, consistent with previous work ^37^. Altogether, these data confirm the specific role of campaniform sensilla and mechanosensory bristles in evaluating stiffness of oviposition substrates.

### Campaniform sensilla and mechanosensory bristles use *painless* as a molecular transducer to gauge surface stiffness

To link our above observations and directly test whether Painless functions in CS and mechanosensory bristle neurons to probe substrate stiffness during egg laying, we set to specifically modulate the expression of *pain* in these neurons. We first downregulated *pain* expression in CS and mechanosensory bristles of the legs, using the same set of Gal4 lines as above to drive a *UAS-pain-IR* construct ^38^, and asked whether the resulting RNAi would phenocopy the effects observed when neurotransmission was blocked in the same neurons. Downregulating *pain* in chemosensory neurons innervating chemosensory bristles in the legs labelled by *poxn-Gal4* only showed a significant effect in oviposition preference with respect to one of the controls (Fig. 4A, Fig. S8A). However, we observed a significant reduction of oviposition preference for the softer substrate when knocking-down *pain* with *CS2-Gal4*, *CS3-Gal4* (Fig. 4B-C, Fig. S8B-C) or *R38B08-Gal4* in the legs (Fig. 4E, Fig. S8E) compared to control flies. Together, these results demonstrate that the protein Painless is directly required in mechanosensory neurons of the leg to mediate egg-laying site selection. The role of *pain* in chemosensory bristles remains, however, unclear. Of note, downregulating *pain* with *CS1* only resulted in a slight and non-significant reduction of preferences for softer substrates, perhaps explained by the different subset of sensory organs targeted by this line (Fig. 4D, Fig. S8D).

**Figure 4.**
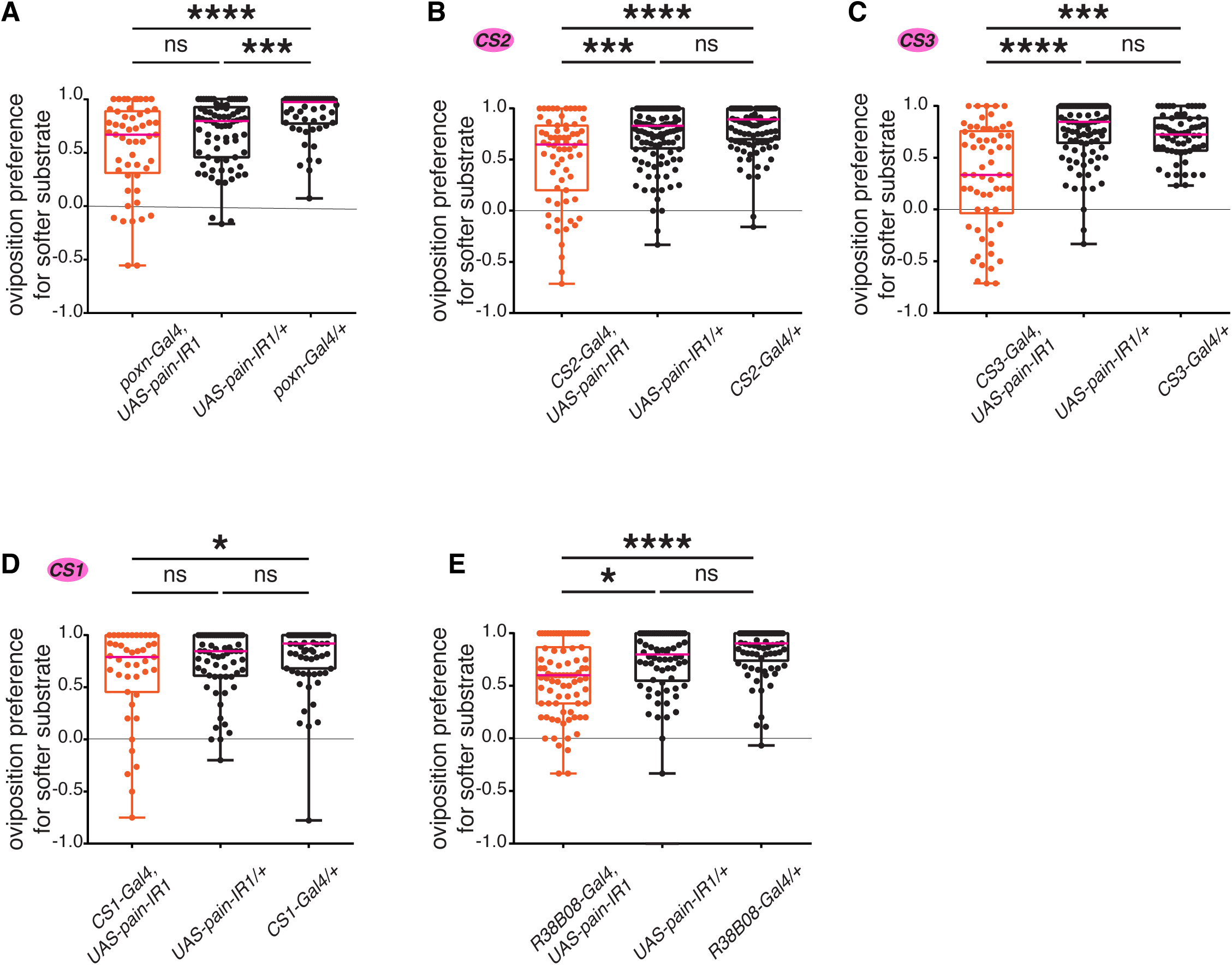
*pain* expression is required in campaniform sensilla and mechanosensory bristles to mediate substrate stiffness. Single-fly two-choice stiffness assay (0.5% vs. 1.25% agarose) of females with RNAi-mediated down-regulation of *pain* expression in neurons innervating chemosensory bristles (A), campaniform sensilla (B-D), or mechanosensory bristles (E). (A) *poxn-Gal4*, *UAS-pain-IR1*, compared to *UAS*-alone or *Gal4*-alone controls (n=53, 76, and 54, respectively). (B) *CS2-Gal4*, *UAS-pain-IR1*, compared to *UAS*-alone or *Gal4*-alone controls (n=67, 120, and 112, respectively). (C) *CS3-Gal4*, *UAS-pain-IR1*, compared to *UAS*-alone or *Gal4*-alone controls (n=62, 102, and 62, respectively). (D) *CS1-Gal4*, *UAS-pain-IR1*, compared to *UAS*-alone or *Gal4*-alone controls (n=43, 66, and 71, respectively). (E) *R38B08-Gal4*, *UAS-pain-IR1*, compared to *UAS*-alone or *Gal4*-alone controls (n= 83, 70, and 77, respectively). p values for statistical significance were calculated using Kruskal–Wallis test followed by a Dunn’s multiple comparisons test. ns, non-significant, p > 0.05; *, p < 0.05; **, p < 0.01; ***, p < 0.005; ****, p < 0.0001. See Figs S8 and S9 for total egg counts.

To strengthen the link, we sought to rescue the stiffness sensing phenotype observed with *pain* mutants, by expressing a *UAS-pain* construct under the control of mechanosensory Gal4 lines. Three Pain isoforms are expressed in flies, at least two of which respond to mechanosensory stimuli very differently ^39^. Lacking information on which isoforms are expressed in different leg mechanosensory neurons, we individually expressed two of these isoforms, Painless^p60^ and Painless^p103^, in a *pain* mutant background. We used the drivers *R38B08-Gal4* and *CS1-Gal4* and tested in two different *pain* mutants whether females expressing these isoforms in specific subsets of leg mechanosensory neurons increased their preferences for softer substrates Specifically, each tested female was homozygous for a *painless* allele (*pain^EP2251^*or *pain^EP2451^*) and contained a Gal4 driver (*R38B08-Gal4* or *CS1-Gal4*) and a *UAS-painless* allele (*UAS-pain^p60^::VFP or UAS-pain^p103^::VFP*; ^39^). We observed a partial (Fig. 5C-D, 5F) to near-complete (Fig. 5E) rescue of the preference for softer substrate in several combinations of these alleles (Fig. 5C-F, Fig. S10C-F), indicating that the function of *pain* is specifically necessary to mediate the preference for softer substrates, both in the mechanosensory bristles and some campaniform sensilla, of the legs. Because the *UAS-pain* isoform are inserted in different genomic locations, they may be expressed at different levels, preventing us to comment on the particular role of *painless^p60^*vs. *painless^p103^* during oviposition.

**Figure 5.**
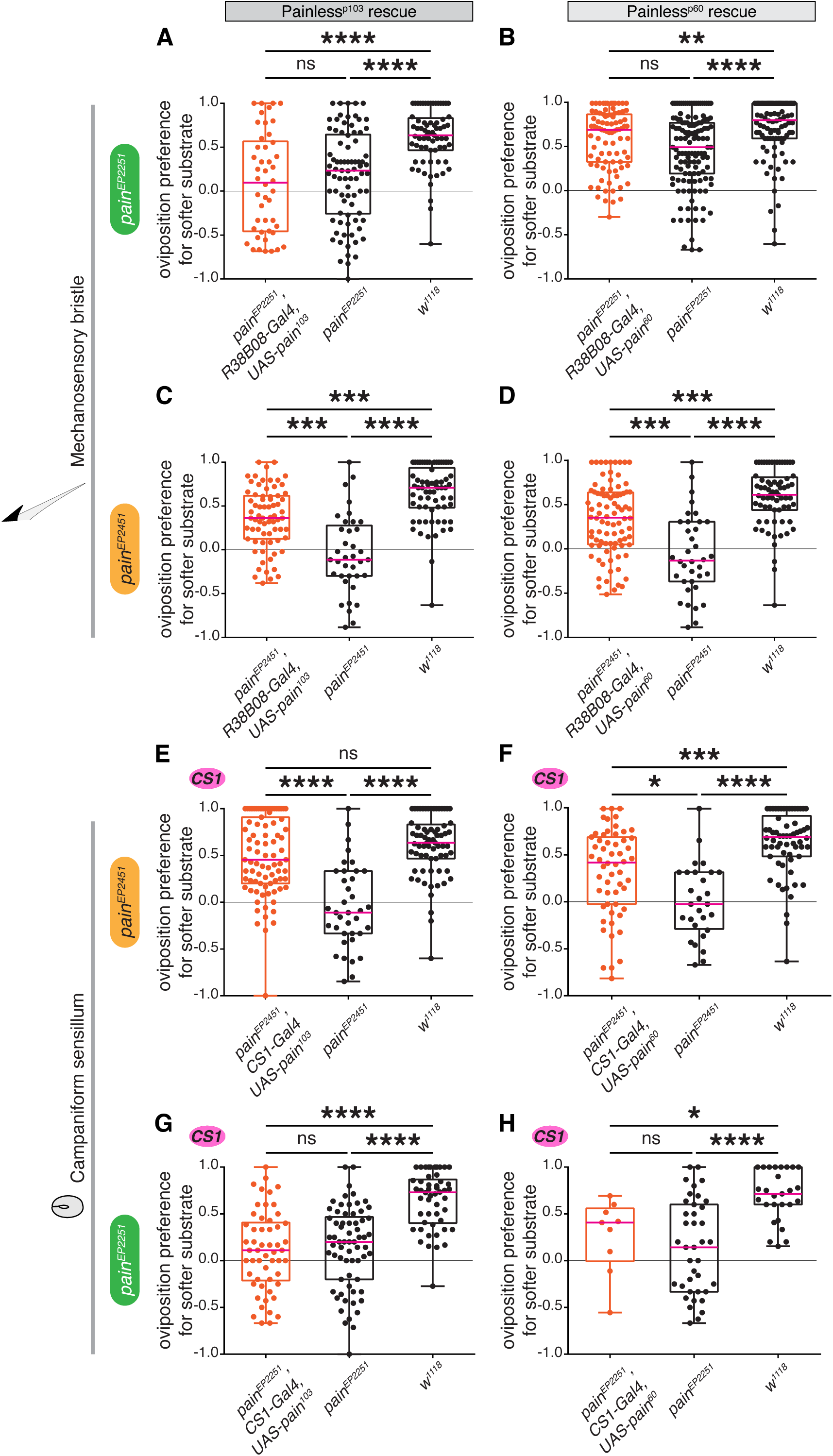
*pain* expression in tarsal mechanosensory organs is sufficient to rescue of *pain* mutants. Overexpression of Painless^p103^ (A, C, E, and G) and Painless^p60^ (B, D, F, and H) isoforms using *UAS-pain^p103^* and *UAS-pain^p103^* in mechanosensory neurons labelled by *R38B08-Gal4* (A, B, C, and D) and *CS1-Gal4* (E, F, G, and H) in *pain* ^[EP2251]^ (A, B, G, and H) and *pain* ^[EP4251]^ (C, D, E, and F) mutant backgrounds (A: n = 46, 82, and 69, respectively; B: n = 87, 127, and 100, respectively; C: n = 71, 37, and 66, respectively; D: n = 88, 36, and 69, respectively; E: n = 83, 37, and 69, respectively; F: n = 56, 28, and 62, respectively; G: n = 54, 44, and 50, respectively; H: n = 9, 39, and 27, respectively). p values for statistical significance were calculated using Kruskal–Wallis test followed by a Dunn’s multiple comparisons test. ns, non-significant, p > 0.05; *, p < 0.05; **, p < 0.01; ***, p < 0.005; ****, p < 0.0001. See Fig. S9 for total egg counts.

Together, our work shows that Painless is necessary to mediate preferences for softer substrates in the context of oviposition, specifically functioning in mechanosensory neurons that innervate subsets of mechanosensory bristles and campaniform sensilla of the tarsi.

## Discussion

Although several pieces of evidence suggest that fruit flies can detect differences in the stiffness of oviposition substrates ^11,12^, the mechanisms behind this ability remain elusive. Expanding on the existing knowledge of a handful of mechanosensory pathways known to influence egg-laying decisions ^12,19^, our work provides evidence for a novel *painless*-dependent mechanosensory pathway in the tarsi that regulates the selection of softer oviposition patches by gravid *D. melanogaster* females. Though the ovipositor might also play a role ^16^, our work only focuses on the contribution of tarsi, as stiffness-sensing peripheral appendages, involved in the initial assessment of stiffness during the search of an amenable oviposition surface. Our findings suggest that the stiffness-sensing function of tarsi stems from the activity of sense organs housed in them, innervated by *pain*-expressing mechanosensory neurons, namely ventrally positioned mechanosensory bristles in the proximal tarsomeres, and campaniform sensilla located dorsally near tarsal joints. We speculate that each type of sensory organ is involved in a distinct way in the stiffness assessment of a plausible surface. The involvement of ventral mechanosensory bristles appears straightforward ^19,35^, as they may directly contact the substrate. We hypothesize that, because the forces opposed by a hard substrate differ from those opposed by a softer substrate, the response of mechanosensory bristles in direct contact with the substrate may be different, with different levels of mechanotransduction at the proximal tarsomeres (where *pain* is expressed) between these conditions.

As for campaniform sensilla, how they influence egg-laying decisions in female flies even though they do not directly contact the substrate, is more elusive. Campaniform sensilla are known to be very sensitive organs ^40,41^ and are seated in the cuticle at joints that experience maximum strain ^42,43^. We propose that CS indirectly sense differences in substrate stiffness by sensing differential compression of the cuticle, in response to the texture of the surface onto which a fly stands. A stiffer substrate exerts higher ground reaction forces ^44–46^ and possibly higher plantar pressure on the tarsi ^47^, increasing strain on the cuticle. Additionally, the rigidity of a stiffer substrate possibly limits force dissipation via substrate deformation ^48^, reflecting these forces back on the tarsi, increasing their load sensing ^49^.

We also show that *pain* function is required in neurons innervating these mechanosensory organs, where it possibly functions as a transducer. Previous studies have indeed shown that in campaniform sensilla of the haltere, the mechanosensitive ion channel *NompC* is a structural component of the mechanical signaling pathway at the distal tips of neurons innervating the cap ^50,51^. A force-induced compression of CS is expected to cause a vertical displacement of the cap, thus stretching the the membrane microtubule connectors at the dendritic tips that gate the NompC channels ^52^. The transmission of mechanical strain to the CS dendrite can lead to the production of receptor potentials with nanometer-scale sensitivity ^53,54^. Similarly, force-induced deflection of mechanosensory bristles also leads to activation of mechanosensory ion channels and subsequent neural action potentials ^55^. Despite not being a force-gated channel itself, Painless expressed in sensory neurons has been previously reported to be involved in several instances of mechanosensation, including in the selection of softer food substrates in *Drosophila* larvae ^56^ and gravity sensing in the antenna of adult flies ^57^. Moreover, other TrpA channels having similar nociceptive functions as Painless have been reported to be required in several mechanosensory behaviors, for instance, nose touch responses and foraging in *C. elegans* ^58^, normal mechanical responsiveness in multiple subtypes of cutaneous sensory neurons in mice ^59^ and sensing shear forces in the intestinal stem cells in *Drosophila* ^60^. In summary, TrpA channels like Painless are also required in the transduction of non-nociceptive mechanical stimuli, potentially functioning as a direct sensor or modulator. In the light of these studies, our data suggests that compromised *painless* activity may disturb the effective registration or transduction of force-related structural alterations in campaniform sensilla and mechanosensory bristles of the legs. A stiff substrate will exert more resistive forces, and thereby cause a stronger dorsal compression of CS and deflection of mechanosensory bristles in the tarsi, thus leading to a different rate of firing and transduction in *pain*-expressing mechanosensory neurons innervating these organs, in comparison to a basal activity when the fly is on a softer patch. Our results are consistent with a recent finding that reports the role of *painless* in the mechanosensory detection of substrate hardness in the context of feeding in fly larvae ^56^.

As for chemosensory bristles, although they may also contact the substrate directly, their role in sensing stiffness is harder to conceive. Despite previous reports on the involvement of chemosensory bristles on the proboscis in sensing textures of feeding substrates ^13,14^, our results do not provide a clear indication regarding their participation in stiffness sensing of oviposition substrates.

We have previously shown that *D. suzukii* differs in the tuning of its sensory systems, correlating with its preference to oviposit on ripening fruits ^11^. These differences further motivated the present study to explore the mechanosensory basis of behavioral divergence between *D. suzukii* and *D. melanogaster*. The correlation between the activity of *painless* and the regulation of oviposition preferences for softer sites in *D. melanogaster*, raises the possibility that changes in *pain* function or expression may mediate the strikingly different egg-laying behaviors between the two species. Previous studies have reported an upregulation of mechanosensory genes such as *pain* in the legs likely yielding heightened mechanosensitivity in *D. suzukii* during ovipositor site selection. By contrast, *D. melanogaster*, has lower expression levels of these mechanosensory genes ^61^. In addition to sensing noxious mechanical stimuli, our results identify a previously unknown substrate-stiffness sensing role of *painless* in the context of oviposition, thus providing a plausible missing connection between differential gene expression of *painless* in *Drosophila* species, and the specific egg-laying behaviors they might influence.

## Acknowledgments

We are grateful to Alexander Blanke, Benjamin Prud’homme, Ilona Grunwald, Lisa Fenk and Anne Kavounoudias for insightful discussions, to Peter Soba, Dan Tracey, Cesar Mendes, Richard Mann, Alexander Borst and Natascha Turetzek for sharing reagents. We thank Richard Benton and Matthieu Cavey for constructive comments on the manuscript. We thank the Bloomington Drosophila Stock Center (NIH P40OD018537) for many fly stocks used in this study and FlyBase ^62^ for genetic and genomic information. This work was supported by the Graduate School of Systemic Neurosciences of Munich (VR) and the Deutsche Forschungsgemeinschaft (GO2495/9-1 to NG).

## Author Contributions

Conceptualization, N.G. and K.C.; methodology and experimental design, N.G., L.B.B., K.C., and V.R.; investigation, V.R., K.C., L.B.B., A.K., C.R., G.F.D., A.P., and K.F.; statistical analysis, V.R.; writing, N.G. and V.R. with input from all authors; supervision, N.G. and A.B.

## Declaration of interests

The authors declare no competing interests.

## Supplemental information

Figures S1-S11

Table S1. *painless* expression in the tarsi

Table S2. p values per figure

Supplemental Files SS1-SS4 (STL). 3D printing files (SS1_base.obj, SS2_teeth.obj, SS3_spacer.obj, and SS4_loft.obj) for various components of the single fly behavioral chamber.

## STAR Methods

### Resource Availability

#### Materials Availability

All requests for fly stocks that have been used/generated in this study are available upon request and will be fulfilled by the Lead Contact, Nicolas Gompel.

#### Data and Code Availability

No codes were generated in this study to analyse behavioral data sets. The preference indices were computed on Microsoft Excel based on the number of eggs laid on each side, followed by application of experiment-specific statistical tests on GraphPad Prism. The graphs were also generated using the same software.

### EXPERIMENTAL MODEL AND SUBJECT DETAILS

Fruit fly, *Drosophila melanogaster* stocks were raised on a standard cornmeal food at 25°C and 45-50% relative humidity on a 12 h light/12 h dark cycle. We used Canton S, *w*^−^, and *w^1118^* as wild types. For detailed transgenic fly stock sources and genotypes, refer to Resources Table. Except for RNAi crosses (at 29°C) and shibire-inactivation experiments (at both 21°C and 31°C), all genetic crosses were set in a 25°C climate chamber. F1 females that were tested in the behavioral assays, had mixed genetic backgrounds. For controls, the UAS and the Gal4 line were crossed with *w^1118^* or *w^−^* to generate F1 heterozygotes.

Transgenic stocks from Bloomington Drosophila Stock Center (BDSC) included *pain[EP2251]* (31432)*, pain[EP2451]* (27895), *pain-Gal4* (27894), *nompA[MB11221]* (27897), *Df(2R)vg135, nompAvg135* (1642), *Piezo[KO]* (28770), *ppk[MI04968]* (38075), *R34E03-Gal4* (48123), *poxn-Gal4* (66685), *R38B08-Gal4* (49541), *VT002828 p65.AD* (73214), *VT039623-GAL4.DBD* (71784), *VT026750-p65.AD* (74539), *VT050230-GAL4.DBD* (71422), *UAS-pain-IR1* (51835), *UAS-pain-IR2* (61299), *40XUAS-IVS-mCD8::GFP* (32195), and *20XUAS-IVS-CsChrimson.mVenus* (55134). Mutants *nan36a,* and *dTrpA1W903* were gifts from Dr. C. Kim (Chonnam National University) and Prof. Dr. Peter Soba (University of Erlangen-Nuremberg) respectively. *10xUAS-Kir2.1-eGFP* and *UAS-shi^ts^* were received from Prof. Dr. Ilona Grunwald Kadow (University of Bonn) and Prof. Dr. Toshihiro Kitamoto (University of Iowa). *Dac[H1]-Flp, 10xUAS>myrtdTomato-SV40>eGFPKir2.1 attp2, UAS-painless-p103:VFP and UAS-painless-p60:VFP, and 6xEGFP,20XUAS* were gifted by Prof. Dr. Richard Mann (Columbia University), Dr. Troy R. Shirangi (Villanova University), Prof. Dr. Dan Tracey (Indiana University), and Dr. Natascha Turetzek respectively.

### METHOD DETAILS

#### Generation of split-Gal4 lines

We targeted specific campaniform sensilla following the strategy described in ^63^. We first selected VT ^64^ and GMR ^65^ enhancer tiles that have expression in the target neuron using the images ^66^ and computational tools ^67–69^ developed in Janelia Research Campus, HHMI. We then paired the corresponding split hemi drivers and screened for combinations that sparsely labelled the target neuron.

#### Behavioral Assays

All the behavioral experiments performed in this study are two-choice oviposition assays that test the oviposition substrate preferences of mated *D. melanogaster* females in various experimental and genetic contexts. Females used in egg-laying experiments were genotyped under CO_2_ anesthesia within 1 d of eclosion and transferred to fresh food vials, allowing them to age for 5-8 days before testing their egg-laying choices.

##### 1. Two-choice-stiffness assay

The oviposition preferences of gravid females for differently stiff substrates were tested using two experimental approaches, group and single-fly assays.

###### *Group-assay* (Panels 1A-E, 2A)

Batches of 100-150 flies containing both males and females were aspirated into the behavioral chamber, comprising a plastic cylinder (14 cm length x 8 cm diameter) covered by a ventilated lid and a base made from a plastic Petri dish (9 cm diameter). The petri dish contained two halves of 0.25% and 0.75% agarose (Biozym 840004) substrates, supplemented with 0.5M glucose (Sigma G8270). 0.75% agarose was first prepared and then poured to the brim of the petri dish. Once the solution solidified, one-half of the substrate was cut out and 0.25% agarose solution was poured into the left-out space. The petri dish containing identical and equally sweetened 0.25% and 0.75% agarose halves represented two substrate choices that differed in substrate stiffness. The two-choice assay was performed for 16 h in a LD cycle (1D; 8 h light and 8h dark) or in complete darkness (1B-1C, 1E, 2A; 16 h in dark), in a climate chamber set at 25°C, 45-50% humidity. Oviposition preference indices (PI) for the softer side (0.25% agarose) were calculated as follows; PI=(#eggs on 0.25% agarose – #eggs on 0.75% agarose)/(#eggs on 0.25% agarose + #eggs on 0.75% agarose).

###### Ablation Experiment (Panels 1B-C)

The proboscis and tarsi of anesthetized adult females were clipped manually with tweezers 3 days before the oviposition assay. The operated flies along with the non-operated ones were subjected to a group-two-choice stiffness assay (40 mated females were used) using the experimental conditions described above.

###### *Single-Fly Assay* (Panels 1F-G, 2B, 3A-H, 4A-E, 5A-H)

In preparation for the assay, relatively smaller fly batches were maintained in food vials in a ratio of 4 females to 5 males, with a minimum of 12 and a maximum of 20 females per vial ^4^, for 5-8 days, starting from the day of eclosion (marking the day of eclosion as Day 1). The food vials containing the flies were placed in an inclined manner to liquefy the media faster. This promotes the retention of eggs in gravid females, forcing them to lay more eggs upon encountering more suitable substrates in the behavioral assay chambers that contain relatively favorable substrates. The behavioral apparatus was assembled using four 3D-printed components: base, teeth, spacer, and loft with a porous acrylic plate. The base has 9 lanes (Fig. S11B), into which about 2.75 mL of 0.5% and 1.25% agarose solutions (supplemented with 3% Acetic Acid (Roth 3738.1) and 0.5M glucose in all single fly stiffness assays except for panels 1F and 2B which only used 3% Acetic Acid as the egg laying stimulant) were poured alternatively. The teeth (Fig. S11A) was then placed on top of the base, creating 56 individual cells, each with two substrate choices. The base and the teeth comprise the lower part of the behavioral chamber). A plastic film was then placed over the teeth which covered its entire surface. The spacer glued to the loft (Fig. S11D-F) were then fitted on top of the mesh to create small cubical spaces to host single mated females. The acrylic plate (Fig. S11G) glued to the surface of the loft has 56 holes, with each hole positioned right on top of a cubical cell (Fig. S11G-H), to facilitate the aspiration of a single fly into a cell. The acrylic lid also has much finer holes to facilitate ventilation in the set-up. The spacer, loft and the acrylic lid make up the upper part of the chamber. Both the parts were held in place with the help of rubber bands on the sides, to assemble the behavioral chamber and gravid females were then aspirated in, followed by covering of the holes with the help of strips of tape to prevent the escape of flies. To start the experiment, the plastic film was removed to allow aspirated flies to touch the substrates. The experiments were initiated ±1.5-2.0 h of lights off, in a climate chamber set at 25 ± 1°C, 45-50% humidity and 12 h light: 12 h dark cycle, allowing exposure to 1.5-2.0 h of homogenous light before lights off and 1.5-2.0 h of complete darkness after lights off. For figure panels 1F, and 2B, the entire experiment ran in darkness over the entire course of 4 h. Oviposition preference indices for the softer side (0.5% agarose) were calculated as follows; PI = (#eggs on 0.5% agarose – #eggs on 1.25% agarose)/(#eggs on 0.5% agarose + #eggs on 1.25% agarose). Only cases where at least five eggs were laid were considered for the calculation of PIs.

###### Incremental Two-Choice-Stiffness Assay (Panel 1F)

Single-fly two-choice assay chambers were used using the same experimental conditions as described above but the agarose concentration was incrementally increased by 0.25% (from 0.5%-2.25%) to constitute the hard substratum while keeping the soft side constant at 0.5% and only 3% acetic acid was used as an egg-laying stimulant. The egg-laying preferences of the females for the softer patch were computed to observe until which agarose concentration pair, a progressive increase in preference was observed. 0.25% agarose evaporates fast, hence were not used in this setup.

###### *RNAi experiments* (Panels 4A-E, S9A-E)

Crosses were performed at 29 ± 1°C, 45-50% humidity, and 12-h light: 12-h dark cycle. Eclosing F1 progeny females were also maintained for aging at 29°C for 5-8 days, starting from the day of eclosion as described above to enhance the efficiency of RNA interference. These flies were then tested for their egg-laying preferences in a single fly two-choice stiffness assay using conditions described above.

###### Shibere inactivation (Panels 3A and 3E)

Crosses were set at 21°C. The eclosing F1 progeny (5-6 days old) were segregated into two batches and tested for their oviposition preferences in a two-choice stiffness assay at 21°C (permissive temperature)) and 31°C (restrictive temperature), to activate the functioning of shibere. The other experimental conditions like humidity and LD cycle remain same as above.

##### 2. Two-choice sugar assays (Panels S6A-D)

Single-fly two-choice assay chambers were used using the same experimental conditions as described above but egg-laying substrates differed in their glucose content instead of stiffness (agarose concentration). The two substrate choices involved plain 1% agarose (without glucose) and 200 mM glucose in 1% agarose. Oviposition preference indices for the sugar containing side were calculated as follows; PI = (#eggs on sugar-containing agarose – #eggs on plain agarose)/(#eggs on sugar containing agarose + #eggs on plain agarose).

#### Imaging

##### Visualization of *pain* in the tarsi (Panels 2D-2E)

For visualizing *painless* expression in the tarsi*, pain-Gal4* was crossed with *20xUAS-6xGFP* line that expresses an enhanced cytoplasmic green florescent protein under the control of the *pain*-Gal4 driver. F1 individuals (6-8 days old females; *pain-Gal4, 20xUAS-6xGFP*) resulting from the cross were anesthetized and whole legs were removed and mounted in DABCO medium (Roth 0718) for visualization. Images were recorded using a Leica SP5-2 point scanning confocal microscope, using a HCX PL APO 63x/1.30 GLYC CORR CS 21 objective. The GFP fluorescence was excited using a 488 nm Argon laser and its fluorescence emission was measured in the spectral region from 530 to 570 nm using a photomultiplier tube detector; this specific range was selected to minimize background autofluorescence. Transmitted light images were also recorded to visualize the external morphology of the legs. The resolution of the images was 2000*800, with dimensions 123*123*504 nm^3^ (x*y*z axes). Resulting images were projected (maximum intensity) using the Fiji ^70^ and merged into composite images.

##### Visualization of the *dac-FLP, UAS>myrtdTomato-SV40>eGFPKir2.1* cassette in the brain and ventral nerve cords (Panels S4A-D’)

Experimental females in which *dac-flippase, UAS>myrtdTomato-SV40>eGFPKir2.1* was expressed under the control of specific drivers, were first anesthetized with CO_2_ and then fixed (2% paraformaldehyde in 75 mM lysine and 37 mM sodium phosphate buffer, pH 7.4) for 2 h at room temperature. The brains, and ventral nerve cords were dissected in PBS containing 0.3% Triton X-100 (PBST), blocked with 10% normal goat serum diluted in PBST for 30 min and incubated in a primary antibody mix overnight at 4 °C, followed by washing in PBST for multiple rounds. The tissues were then incubated in a secondary antibody mix overnight at 4 °C. A final round of PBST washing occurred before the tissue was mounted using Vectashield (Vector Laboratories, RRID:AB_2336789) and visualized using an LSM 710 laser-scanning confocal microscope with a ×25/0.8 DIC or ×40/1.2 W objective (Zeiss). Primary antibodies used were mouse anti-Bruchpilot (nc82; 1:10; Developmental Studies Hybridoma Bank), chicken anti-GFP (1:1,000; Aves Labs), and rabbit anti-DsRed (1:500; Clontech). Secondary antibodies used were Alexa Fluor 633 goat anti-mouse, Alexa Fluor 488 goat anti-chicken, Alexa Fluor 488 goat anti-rabbit and Alexa Fluor 555 goat anti-rabbit (all at 1:200; Life Technologies). Acquired images were processed using the Fiji distribution of ImageJ (NIH).

##### Visualizing expression of CS-Gal4s in the brain and ventral nerve cord

To validate the specificity of CS split-Gal4 drivers in labelling only leg campaniform sensilla, the expression of *CS2-Gal4* and *CS3-Gal4*>*20XUAS-CsChrimson-mVenus* females were visualized in the brains and ventral nerve cords (Panels S4E-F’), after processing the tissue samples according to FlyLight protocols (https://www.janelia.org/project-team/flylight/protocols).

##### Visualizing expression of CS-Gal4s in the legs

For identifying CS-expressing drivers, respective drivers were crossed with *UAS-mcD8-GFP* (Panels S3B-K). Whole flies were fixed in 4% paraformaldehyde for 45 minutes, washed four times with PBS and then the legs were dissected and mounted in Vectashield. For visualization (Leica CLSM SP8; Leica Camera AG, Wetzlar, Germany), the 63x water objective was used for generating detailed images and 10x dry objective for the general images. Line average was usually 4, the images were taken in 1024×1024 resolution. The Z-stack differed between the scans. To visualize the cuticle, its autofluorescence in the far-red spectrum was used. Acquired images were processed using Fiji.

### STATISTICS ANALYSIS

GraphPad Prism 6 software (GraphPad Software, San Diego, CA) was used to produce graphs and statistically analyze data. The data from the behavioral assays have been presented using box and whisker plots. These plots depict the median of computed PIs within the box and the whiskers extend to the true minimum and maximum values for each genotype. No sample-size estimation was performed in this study. In group assays, n depicts biological replicates, involving groups of flies of a particular genotype, tested on different days. In single fly assays, n denotes number of individual gravid females tested. For testing statistical significance, inter-genotypes comparisons of oviposition preferences were performed using a Mann Whitney test (Non-Parametric t-Test). When there were more than two groups under comparison, oviposition preference indices were compared using a Kruskal-Wallis test, followed by a Dunn’s Multiple Comparison Test (data sets were not normally distributed, therefore the choice of these specific tests). For all statistical analyses, notations are as follows: ns, non-significant; p > 0.05; *, p < 0.05; **, p < 0.01; ***, p < 0.005; ****, p < 0.0001.

---

**Figure S1.**
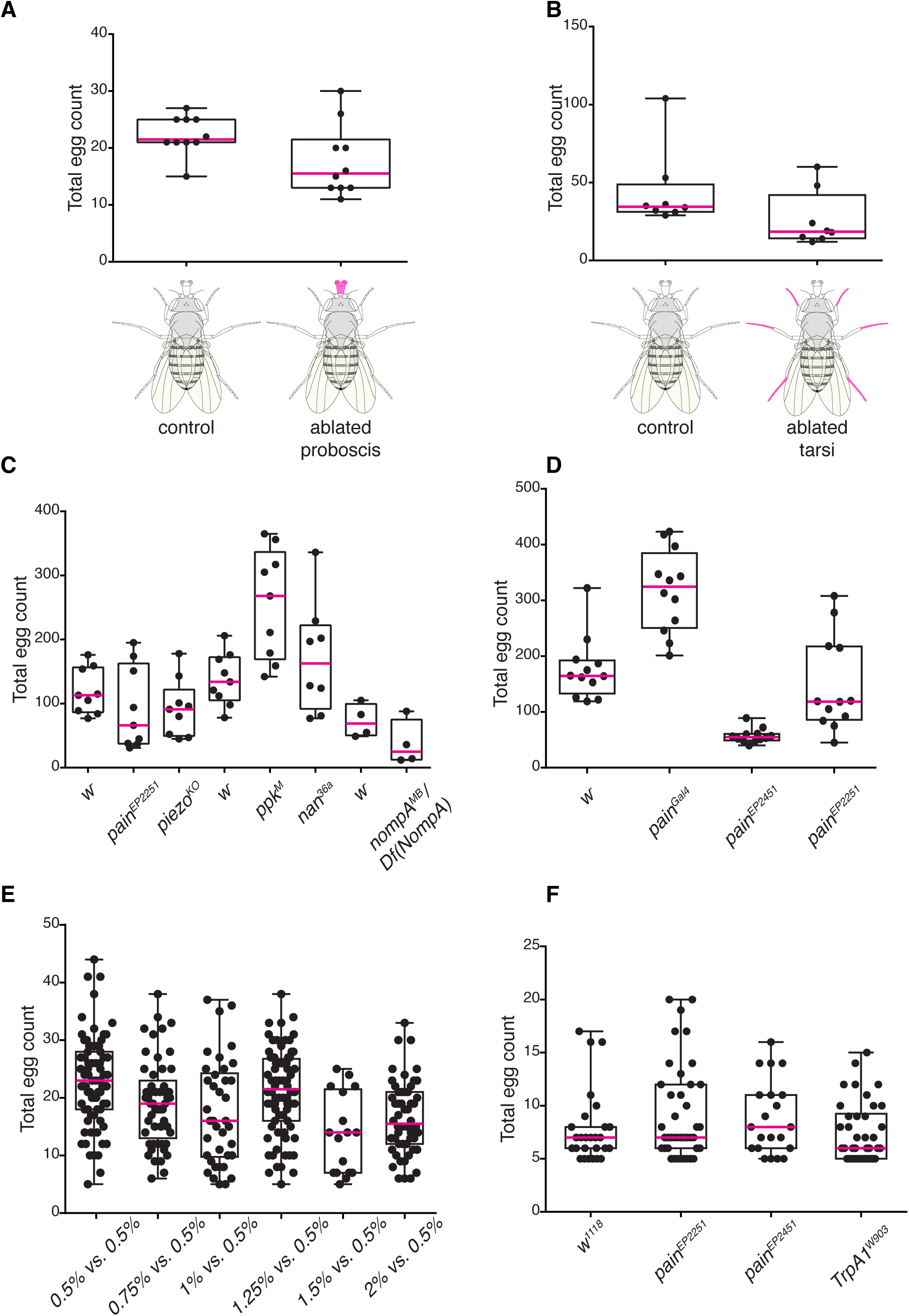
Absolute egg counts used to calculate the preference indices of Figure 1. Relates to Figure 1. (A-F) Total number of eggs laid per replicate during the group assays involving ablation of proboscis (A) or tarsi (B), as well as genetic screening of potential mechanoreceptors (C) and *painless* alleles (D). (E-F) Total number of eggs laid per mated female during the single fly assays testing oviposition preferences of wild-type flies involving increasing agarose concentration (E) and mutant alleles of TrpA channels (F).

**Figure S2.**
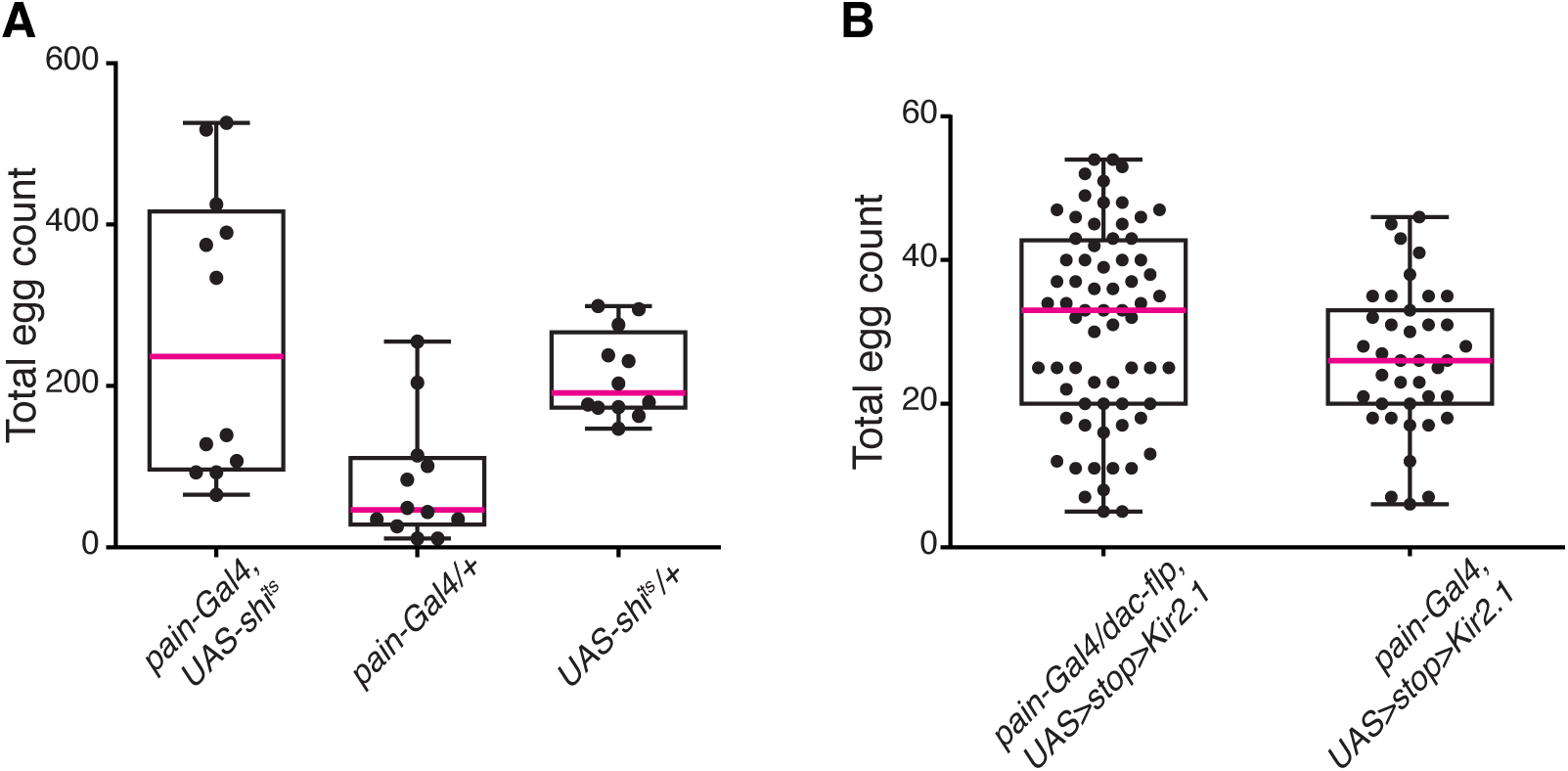
Absolute egg counts used to calculate the preference indices of Figure 2. Relates to Figure 2. (A-B) Total number of eggs laid per replicate during the group assays, upon silencing of all *pain*-expressing neurons labelled by *pain-Gal4* (A) and per mated female during the single fly assays, involving leg-specific silencing of *pain*-expressing sensory neurons using *dac-flippase* (B).

**Figure S3.**
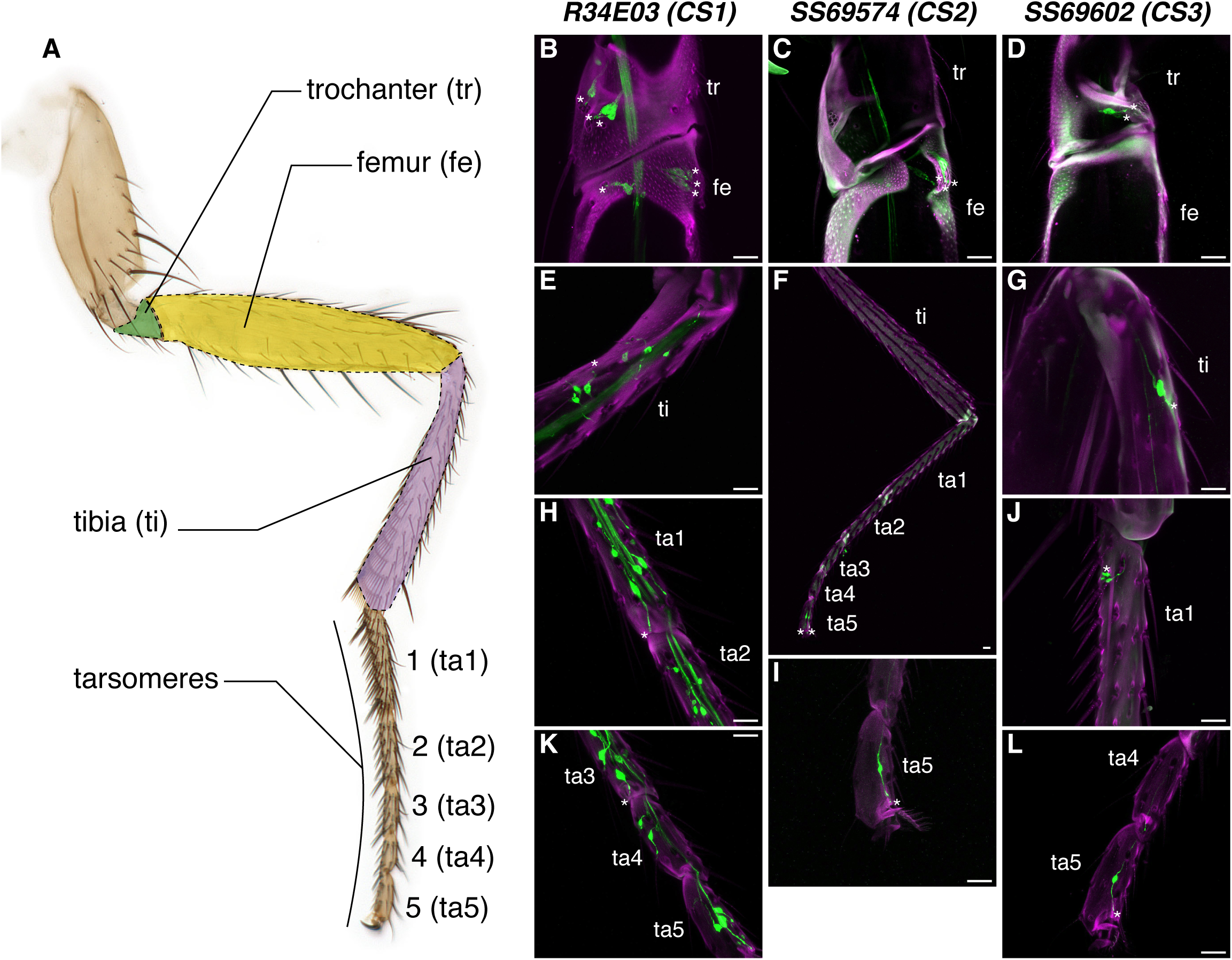
Expression patterns of *CS-Gal4* lines across the mesothoracic leg. (A) Anatomy of a mesothoracic leg. Confocal images of the expression patterns of *CS1-Gal4* (B,E,H,K), *CS2-Gal4* (C,F,I) and *CS3-Gal4* (D,G,J, L) across the mesothoracic leg. The CS labelled by these driver lines are marked with an asterix (*). Scale bar, 20 µm.

**Figure S4.**
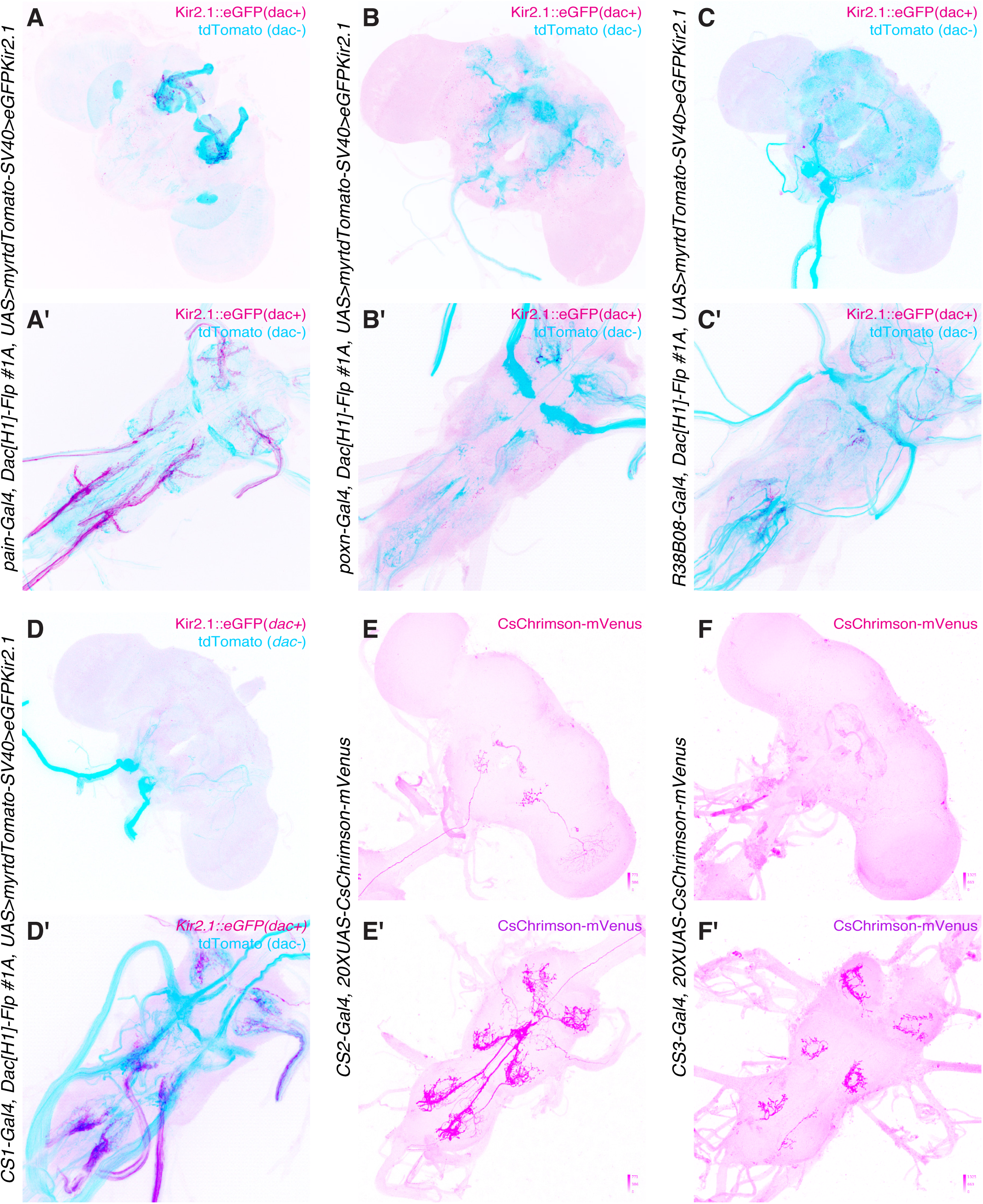
Expression of Gal4 drivers in the central nervous system. Antibody staining against GFP (Kir2.1-eGFP-expressing *Gal4* neurons, magenta) and DsRed (tdTomato-expressing *Gal4* neurons, blue) on brains (A-F) and ventral nerve cord (A’-F’) from females with the following genotypes: (A, A’) *pain-Gal4, Dac[H1]-Flp, UAS>myrtdTomato-SV40>eGFPKir2.1*. (B, B’) *poxn-Gal4, Dac[H1]-Flp, UAS>myrtdTomato-SV40>eGFPKir2.1*. (C, C’) *R38B08-Gal4, Dac[H1]-Flp, UAS>myrtdTomato-SV40>eGFPKir2.1*. (D, D’) *CS1-Gal4, Dac[H1]-Flp, UAS>myrtdTomato-SV40>eGFPKir2.1*. (E, E’) *CS2-Gal4>20XUAS-CsChrimson-mVenus*. (F, F’) *CS3-Gal4>20XUAS-CsChrimson-mVenus*.

**Figure S5.**
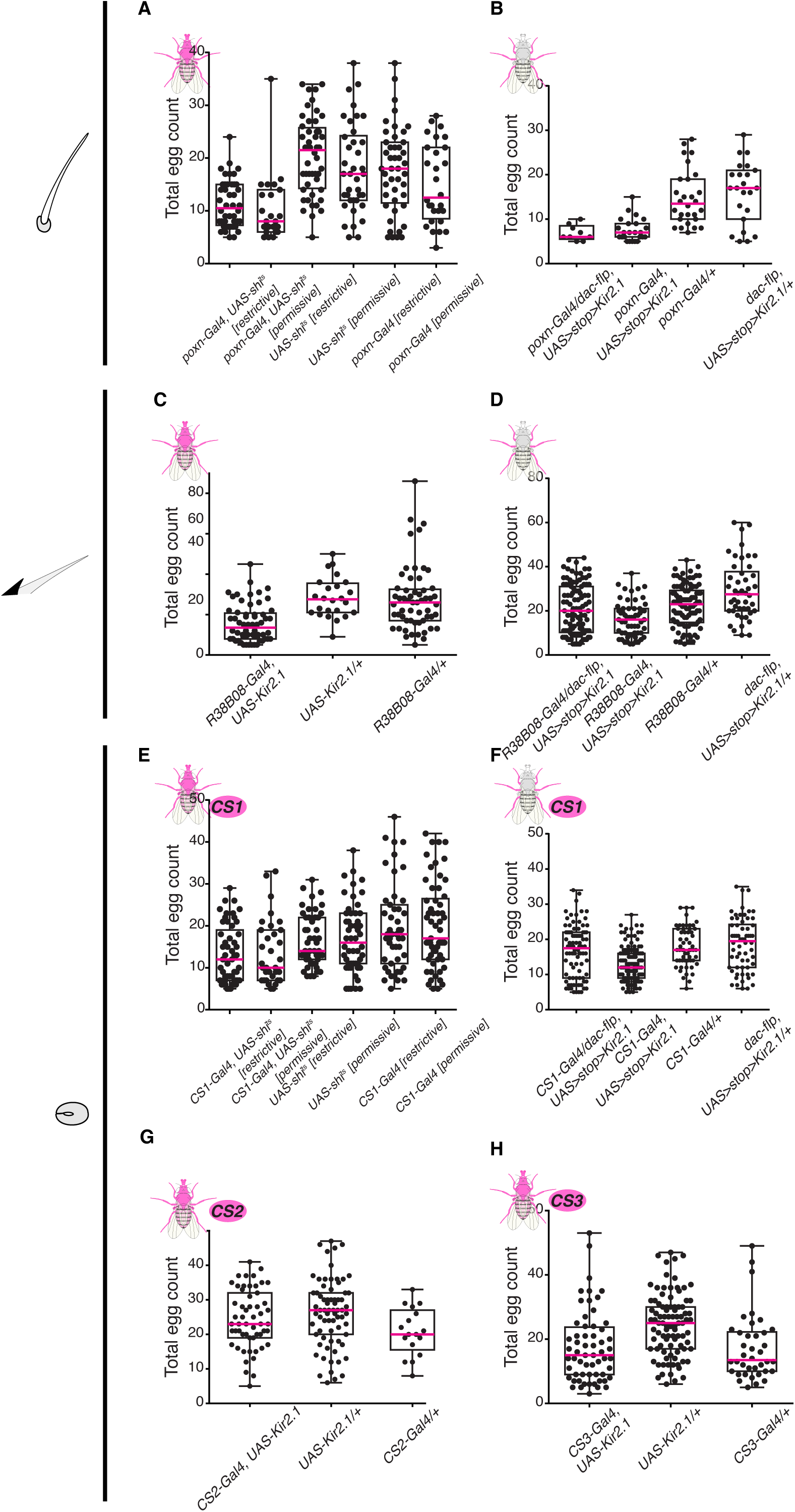
Absolute egg counts used to calculate the preference indices of Figure 3. Relates to Figure 3. (A-H) Total number of eggs laid per mated female, involving Kir2.1- or Shibire^ts^-mediated silencing of neurons labelled by *poxn-Gal4* (A-B)*, R38B08-Gal4* (C-D) and *CS1-Gal4* (E-F), *CS2-Gal4* (G), and *CS3-Gal4* (H).

**Figure S6.**
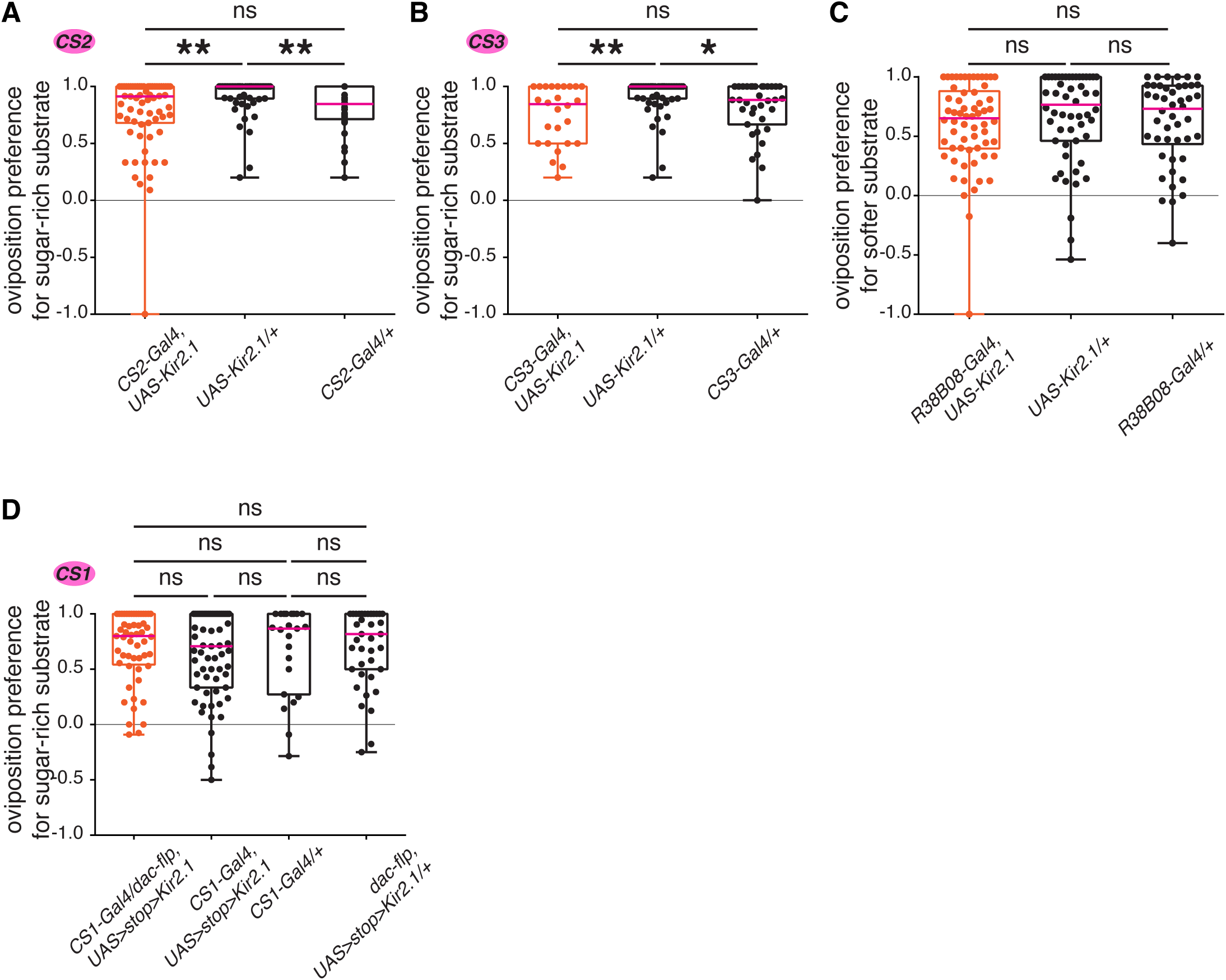
Blocking activity of mechanosensory neurons in sugar preference assays. Single-fly two-choice sugar assay (200 mM glucose + 1% agarose vs. 1% agarose) of females with global (A-C) or leg-restricted (D) blocking of neurotransmission in mechanosensory neurons innervating campaniform sensilla (A-B, D) or mechanosensory bristles (C). (A) *CS2-Gal4*, *UAS-Kir2.1*, compared to *UAS*-alone or *Gal4*-alone controls (n=67, 53, and 33, respectively). (B) *CS3-Gal4*, *UAS-Kir2.1*, compared to *UAS*-alone or *Gal4*-alone controls (n=26, 53, and 35, respectively). (C) *R38B08-Gal4*, *UAS-Kir2.1*, compared to *UAS*-alone or *Gal4*-alone controls (n=62, 54, and 46, respectively). (D) conditional inactivation of *CS1-Gal4* neurons in the legs, (*CS1-Gal4*, *dac-flippase*, *UAS>stop>Kir2.1*), compared to controls lacking one of the genetic components (n= 53, 63, 23, and 39, respectively). p values for statistical significance were calculated using Kruskal–Wallis test followed by a Dunn’s multiple comparisons test. ns, non-significant, p > 0.05; *, p < 0.05; **, p < 0.01; ***, p < 0.005; ****, p < 0.0001.

**Figure S7.**
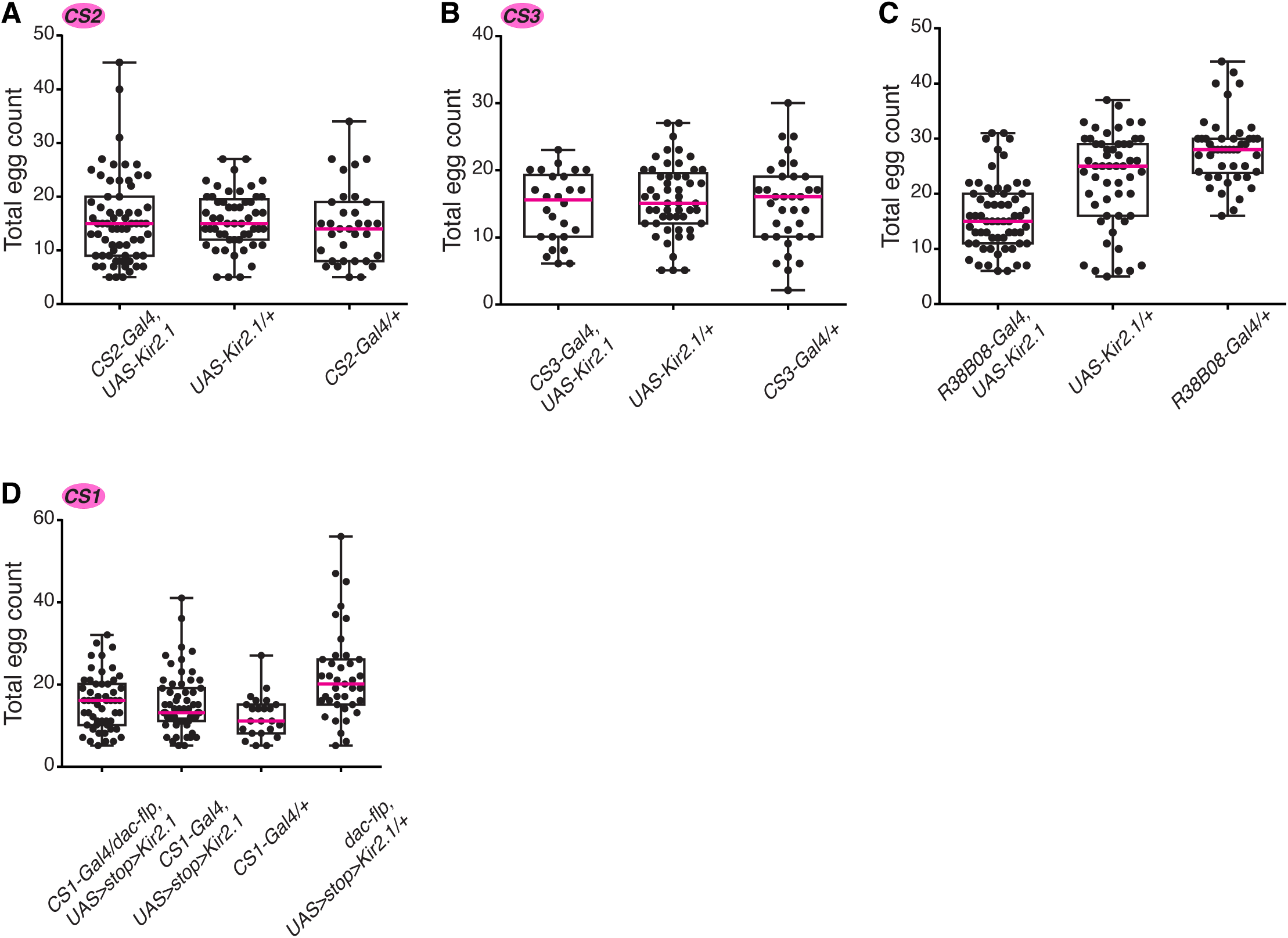
Absolute egg counts used to calculate the preference indices of Figure S6. Relates to Figure S6. (A-C) Total number of eggs laid per mated female, involving Kir2.1-mediated silencing of neurons labelled by *CS2-Gal4* (A), *CS3-Gal4* (B) and *R38B08-Gal4* (C), respectively. (D) Total number of eggs laid per mated female, involving conditional silencing of mechanosensory neurons innervating subsets of campaniform sensilla and mechanosensory bristles labelled by *CS1-Gal4*.

**Figure S8.**
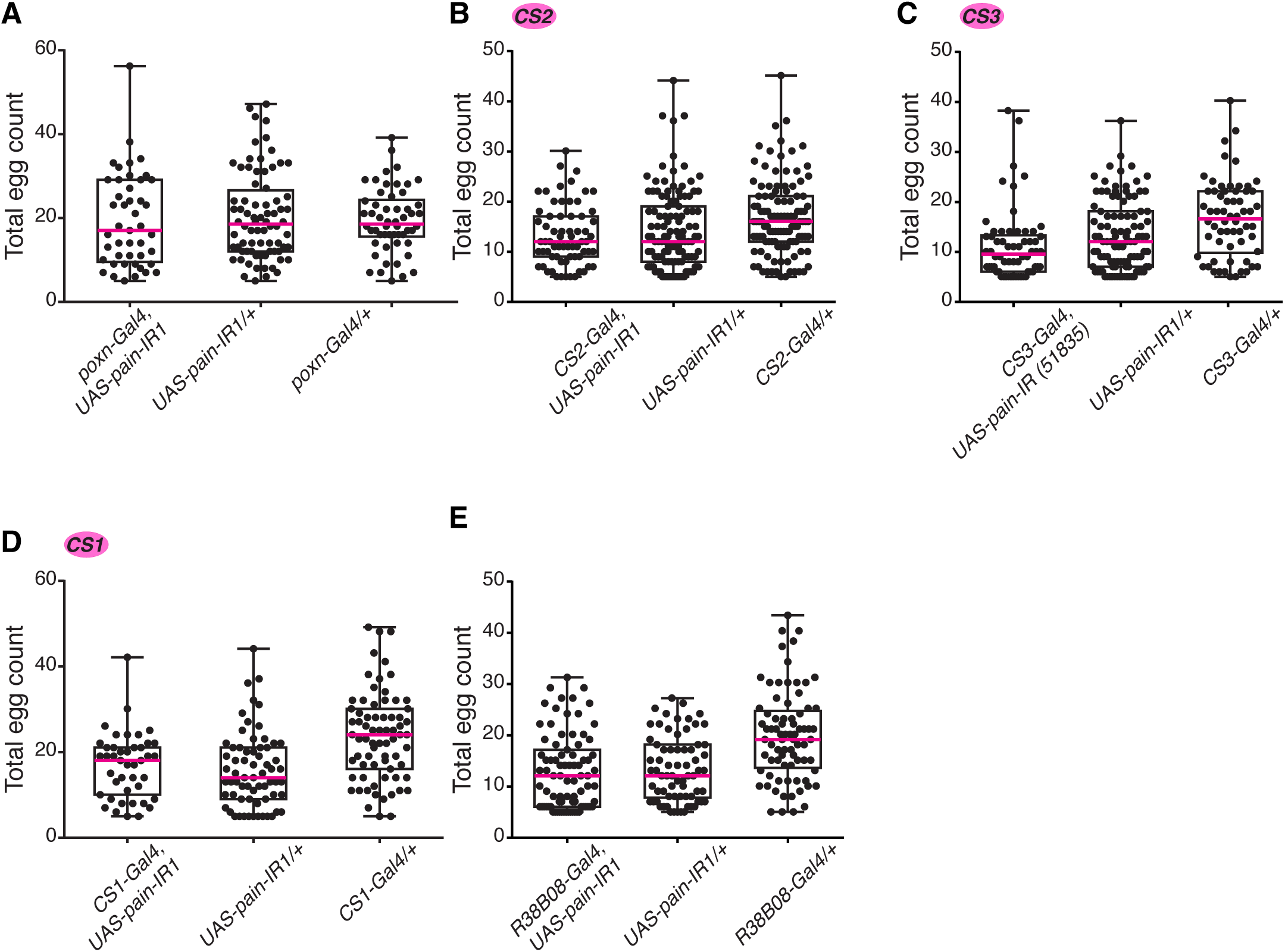
Absolute egg counts used to calculate the preference indices of Figure 4. Relates to Figure 4. (A-E) Total number of eggs laid per single mated female upon knocking down *pain* expression (*UAS-pain-IR1*) in chemosensory neurons associated with chemosensory bristles labelled by *poxn-Gal4* (A), mechanosensory neurons innervating subsets of campaniform sensilla labelled by *CS2-Gal4* and *CS3-Gal4* (B-D), both campaniform sensilla and mechanosensory bristles labelled by *CS1-Gal4,* and only mechanosensory bristles by *R38B08-Gal4* (E), respectively.

**Figure S9.**
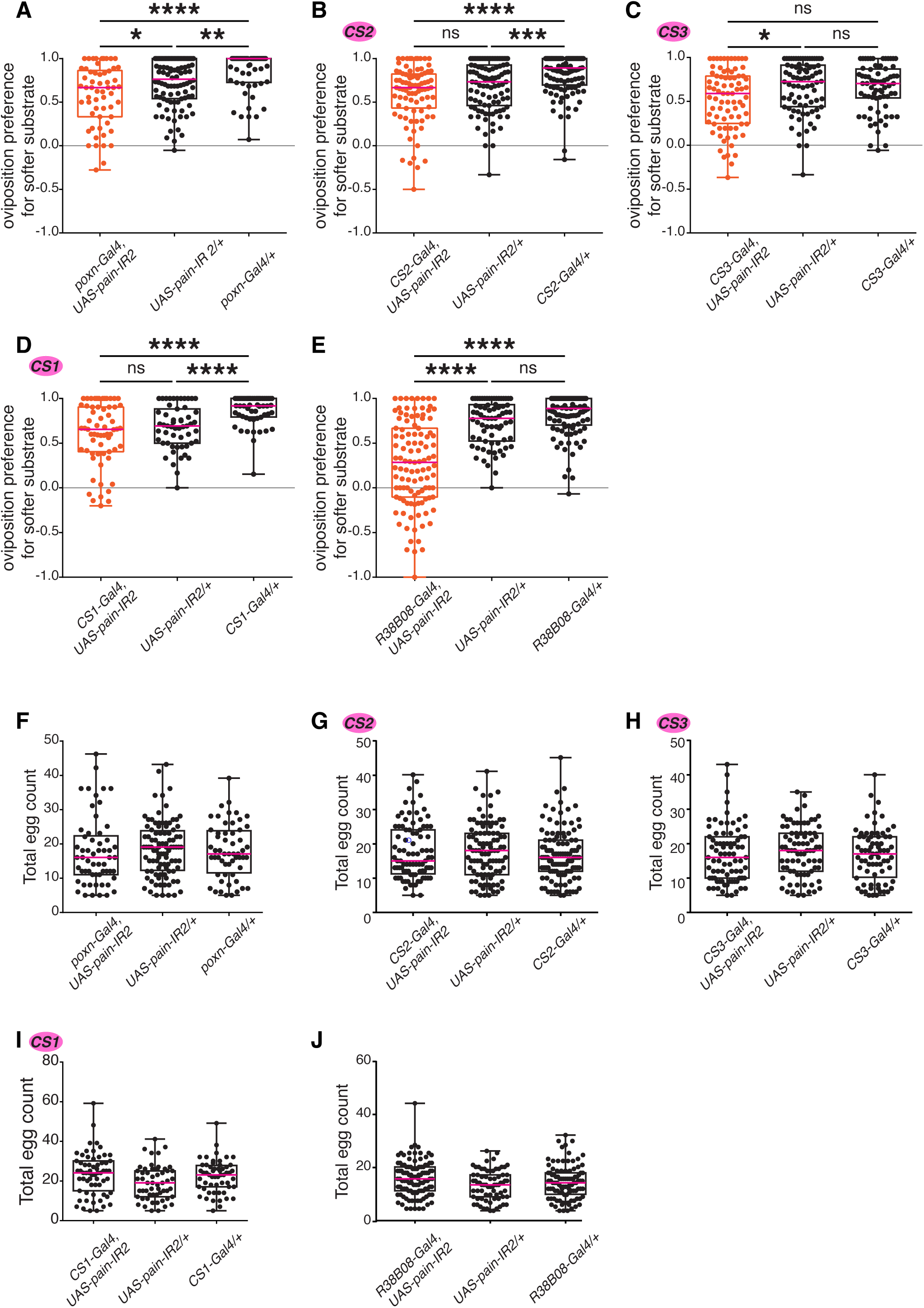
Knocking down *pain* expression using an alternate *UAS-pain-IR2* line. Single-fly two-choice stiffness assay (0.5% vs. 1.25% agarose) of females with RNAi-mediated down-regulation of *pain* expression in neurons innervating chemosensory bristles (A, F), campaniform sensilla (B-D, G-I), or mechanosensory bristles (E, J). (A) *poxn-Gal4*, *UAS-pain-IR2*, compared to *UAS*-alone or *Gal4*-alone controls (n=58, 96, and 56, respectively). (B) *CS2-Gal4*, *UAS-pain-IR2*, compared to *UAS*-alone or *Gal4*-alone controls (n=88, 99, and 112, respectively). (C) *CS3-Gal4*, *UAS-pain-IR2*, compared to *UAS*-alone or *Gal4*-alone controls (n=80, 80, and 72, respectively). (D) *CS1-Gal4*, *UAS-pain-IR2*, compared to *UAS*-alone or *Gal4*-alone controls (n=63, 55, and 56, respectively). (E) *R38B08-Gal4*, *UAS-pain-IR2*, compared to *UAS*-alone or *Gal4*-alone controls (n= 83, 70, and 77, respectively). (F-J) Total egg counts used to calculate preference indices in panels A-E. p values for statistical significance were calculated using Kruskal–Wallis test followed by a Dunn’s multiple comparisons test. ns, non-significant, p > 0.05; *, p < 0.05; **, p < 0.01; ***, p < 0.005; ****, p < 0.0001.

**Figure S10.**
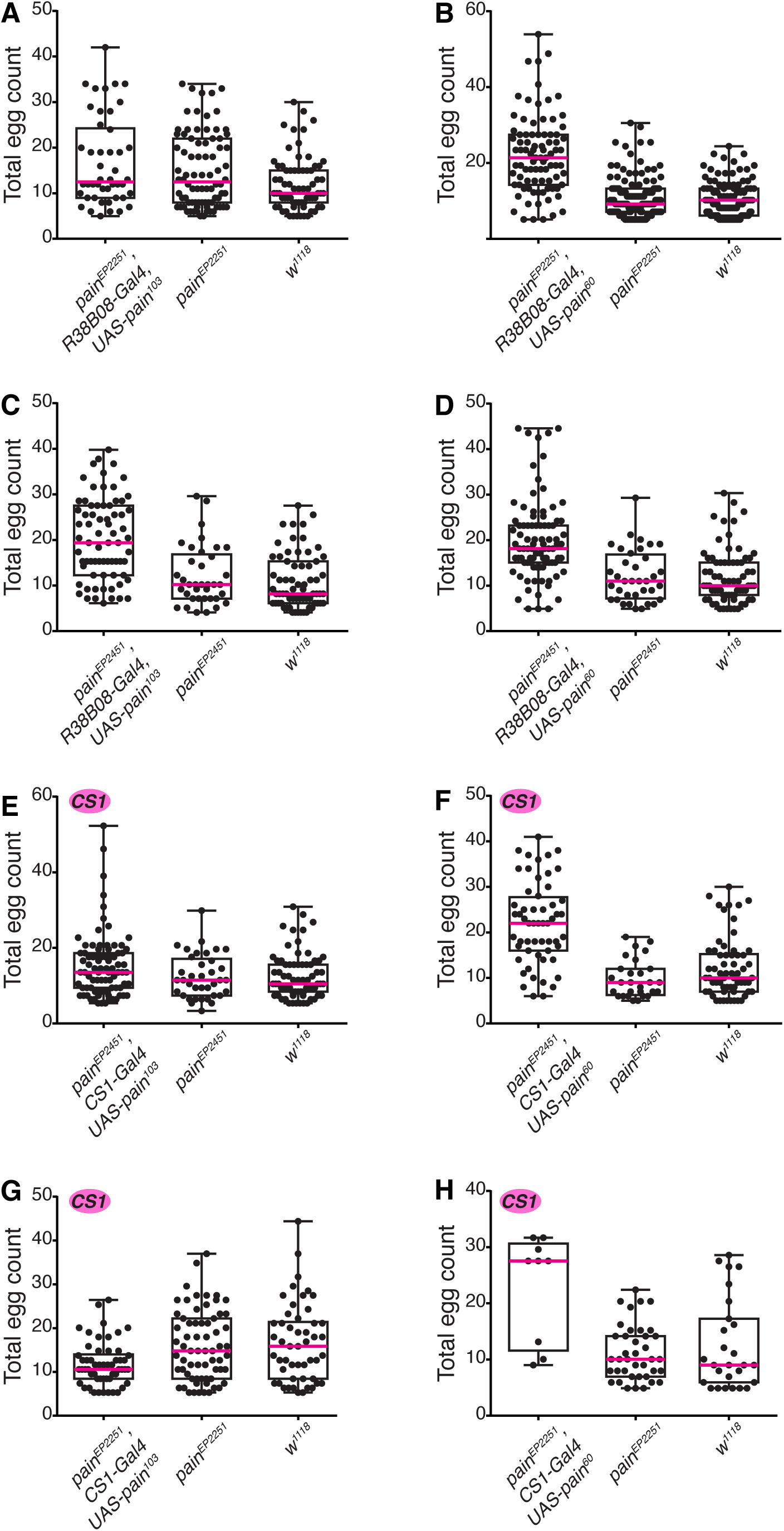
Absolute egg counts used to calculate the preference indices of Figure 5. Relates to Figure 5. Total number of eggs laid upon overexpressing Painless^p103^ (A, C, E, and G) and Painless^p60^ (B, D, F, and H) isoforms using *R38B08-Gal4* (A, B, C, and D), and *CS1-Gal4* (E, F, G, and H) in *pain* ^[EP2251]^ (A, B, G, and H) and *pain* ^[EP4251]^ (C, D, E, and F) mutant backgrounds.

**Figure S11.**
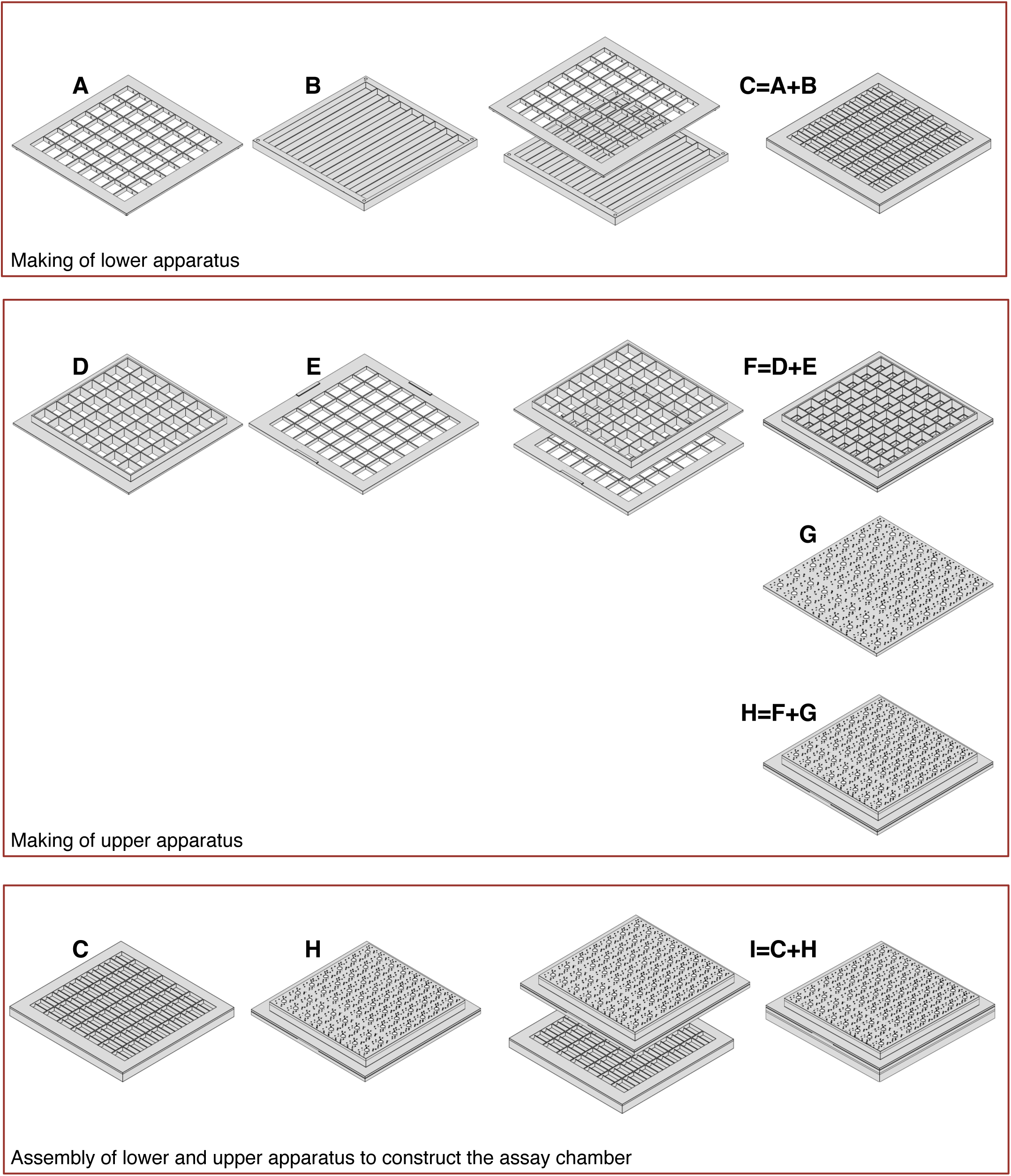
3D model of behavior chambers for 2-choice assay. Assembly of a single fly two-choice oviposition chamber, consisting of a lower and upper half. The divider (A) with walls of 0.5 mm is fitted on top of the base (B) containing 9 lanes into which alternatingly soft and hard substrates are poured, to create the lower half (C). This composite part provides 56 chambers, each containing patches of soft and hard substrates, separated by the divider. To construct the upper half, a spacer (E) is attached to the base of the loft (D), forming a composite unit (F). A porous lid cut out of 1.5 mm acrylic, with 2.8 mm diameter (circular loading ports) is then glued on top of it (G) to constitute the upper half of the behavioral apparatus (H). This upper half (H) is then placed on top of (C), and the entire apparatus is held in place with rubber bands. The final dimensions of an individual chamber is 8.4mm × 10mm (width (x) by length (y)) with a height of 8.4 mm (1 mm (base + divider) + 7.4 mm (spacer + loft).

## Notes

### Competing Interest Statement

The authors have declared no competing interest.

